# Comparison of HAV and HCV infections *in vivo* and *in vitro* reveals distinct patterns of innate immune evasion and activation

**DOI:** 10.1101/2022.11.07.515428

**Authors:** Ombretta Colasanti, Rani Burm, Hao-En Huang, Tobias Riedl, Jannik Traut, Nadine Gillich, Teng-Feng Li, Laura Corneillie, Suzanne Faure-Dupuy, Oliver Grünvogel, Danijela Heide, Ji-Young Lee, Cong Si Tran, Uta Merle, Maria Chironna, Florian F.W. Vondran, Katharina Esser-Nobis, Marco Binder, Ralf Bartenschlager, Mathias Heikenwälder, Philip Meuleman, Volker Lohmann

## Abstract

**Objective:** Hepatitis A virus (HAV) infections are considered not to trigger an innate immune response *in vivo*, in contrast to hepatitis C virus (HCV). This lack of immune induction has been imputed to strong immune counteraction by HAV proteases 3CD and 3ABC. We aimed at elucidating the mechanisms of innate immune induction and counteraction by HAV and HCV *in vivo* and *in vitro*.

**Design:** uPA-SCID mice with humanized liver were infected with HAV and HCV. Hepatic cell culture models were used to assess HAV and HCV sensing by TLR3 and RIG-I/MDA5, respectively. Cleavage of the adaptor proteins TRIF and MAVS was analyzed by transient and stable expression of HAV and HCV proteases and virus infection.

**Results:** We detected similar levels of Interferon stimulated genes (ISGs) induction in hepatocytes of HAV and HCV infected human liver chimeric mice. In cell culture, HAV induced ISGs exclusively upon sensing by MDA5 and dependent on LGP2. TRIF and MAVS were only partially cleaved by HAV 3ABC and 3CD, not sufficiently to abrogate signalling. In contrast, HCV NS3-4A efficiently degraded MAVS, as previously reported, whereas TRIF was not cleaved.

**Conclusions:** HAV induces an innate immune response in hepatocytes via MDA5/LGP2, with limited control of both pathways by proteolytic cleavage. HCV activates TLR3 and lacks TRIF cleavage, suggesting that this pathway mainly contributes to HCV induced antiviral response in hepatocytes. Our results shed new light on induction and counteraction of innate immunity by HAV and HCV and their potential contribution to clearance and persistence.

**SIGNIFICANCE OF THIS STUDY:** *What is already known on this topic?:*

— Despite sharing biological and molecular similarities, HAV infections are always cleared while HCV infections persist in most cases.
— In infected chimpanzees HAV does not trigger a strong innate immune response, as opposed to HCV. This has been imputed to the action of HAV proteases abrogating the signalling pathways.
— Physiological *in vitro* and *in vivo* models, based on human hepatocytes, to assess HAV and HCV mechanisms of induction and interference of innate immunity are still missing.

*What this study adds:*

— HAV induces an innate immune response *in vitro* and *in vivo*, in systems with intact signalling pathways and devoid of adaptive immunity.
— HAV 3ABC and 3CD proteases do not abolish the host innate immune response.
— HCV NS3-4A protease disrupts the RLRs pathways, but cannot cleave TRIF and has no impact on TLR3 response.

*How this study might affect research, practice or policy:*

— This study offers a comprehensive, side-by-side investigation on HAV and HCV infections in physiological models which recapitulate a cytokine response in the human liver, and allows a precise assessment of the viral interference related to the function of the respective signalling pathways.
— Our results elucidate mechanisms, so far controversial or poorly investigated, thus contributing to our understanding of HAV clearance and HCV persistence.

## INTRODUCTION

Hepatitis A Virus (HAV) and Hepatitis C Virus (HCV) are both hepatotropic, (+)-sense RNA viruses marked by similar molecular features, but causing remarkably different infection outcomes. While HAV infections always result in a self-resolving acute infection, HCV infection mainly evolves to chronic, persistent hepatitis. HCV affects >58 million people worldwide, with approximately 290,000 annual deaths^1^ whereas for HAV around 158 million cases and 3900 deaths were reported in 2019^2^. The reasons underlying these strikingly opposed outcomes are poorly understood.

The replication cycles of HCV and HAV share distinct similarities, characterized by the formation of cytoplasmic membranous replicase complexes^3, 4^, and the synthesis of a double-stranded (ds) RNA intermediate which can potentially trigger the cytosolic RIG-I-like receptors (RLRs) Retinoic Acid-Inducible gene I (RIG-I) and Melanoma Differentiation-Associated protein 5 (MDA5) (reviewed in ^5^), both sensing cytosolic dsRNA. The HAV genome is flanked by a 3’ poly(A) tail and a 5’ terminal covalently bound peptide (VPg) (reviewed in ^6^). The genome of HCV contains a free 5’ triphosphate at its 5’ end and a highly conserved structured sequence at the 3’end (reviewed in ^7^). RIG-I, which typically senses dsRNAs with free 5’ triphosphate, is one of the cytosolic sensors for HCV, together with MDA5^8-10^ which is also regarded as the pattern recognition receptor (PRR) responsible for detection of HAV^11^. Both receptors recruit the adaptor protein Mitochondrial antiviral-signaling protein (MAVS) (reviewed in ^5^) and activate a signaling cascade which culminates in the establishment of an antiviral state, based on Interferon (IFN) production and expression of IFN Stimulated Genes (ISGs). Furthermore, the MDA5-triggered IFN response to several (+)RNA viruses was described to depend on Laboratory of Genetics and Physiology 2 (LGP2)^12^. Moreover, HCV dsRNA was reported to trigger the endosomal PRR Toll-Like-Receptor 3 (TLR3)^13, 14^, which also activates IFN responses through recruitment of the adaptor protein TIR-domain-containing adapter-inducing interferon-β (TRIF) (reviewed in ^5^).

Previous studies identified various interference mechanisms counteracting the establishment of an antiviral state. HCV NS3-4A and HAV 3ABC proteases were shown to disrupt RLRs through MAVS cleavage^15-18^. Degradation of TRIF by HAV 3CD was reported^19^ but remains controversial for HCV NS3-4A^20, 21^. Alternative mechanisms of innate immune interference, involving different HAV proteins, were described as well^22-25^. In case of HCV, secretion of dsRNA was furthermore found to weaken TLR3 responses^14^.

Particularly for HAV, only limited sets of *in vivo* data are available. A study investigating intrahepatic innate immune response in HAV and HCV infected chimpanzees found very little ISG induction in case of HAV, whereas HCV infection was associated to robust ISG expression^26^, comparable to what is found in the majority of chronic HCV patients^27, 28^. Yet, chemokines are robustly increased in the serum of acute hepatitis A (AHA) patients^29, 30^. Also, in the murine model productive HAV infection requires the absence of MAVS or type I IFN receptors^31^, suggesting a robust innate immune activation in this model, which cannot be overcome by HAV interference mechanisms^32^. Overall, these data indicate an important contribution of innate immunity to persistence and clearance of HCV and HAV infections. However, so far it remains unclear to which extent the various counteraction mechanisms reported in literature contribute to infection outcome.

Here, we aimed at comprehensively clarifying how HAV and HCV trigger and counteract innate immunity in physiologically relevant *in vivo* and *in vitro* models. We show that HAV and HCV induce to a similar extent ISGs and IFNs in the liver of infected uPA-SCID mice with humanized liver, demonstrating that both viruses trigger innate immunity in the infected human hepatocytes. HAV sensing seems to be MDA5 specific, but requires expression of LGP2. In In line with these data, MAVS cleavage by HAV is only limited. In contrast, HCV sensing by RIG-I and MDA5 is completely blocked by full cleavage of MAVS, as shown before; therefore, the ISG response induced by HCV in hepatocytes appears to primarily originate from TLR3. Consistently, we were not able to detect TRIF cleavage by HCV NS3-4A.

## MATERIAL AND METHODS

### Cells

Huh7-Lunet cells and Huh7-Lunet T7 cells have been described before^33^, as well as the Con1 subgenomic replicon clone 9-13^34^ (gt1b, GenBank accession number AJ238799) and LucubineoJFH1^35^ (gt2a, GenBank accession number AB047639), along with the HAV replicon cell lines^36^. Huh 7.5 cells were a generous gift from Charles Rice. Huh 7.5 TLR3, RIG-I and MDA5 were generated in a previous study^14^. The HepaRG LGP2 KO cell line cell line was described recently^37^. HepG2 were kindly provided by H. P. Dienes. RIG-I KO and MDA5 KO cells were generated by CRISPR/Cas9 in this study. All cell lines used in this study were validated and free of mycoplasma. PHH were isolated from liver resections of patients undergoing partial hepatectomy as described previously^14^.

### Viruses

For HCV infection of Huh7.5 cells, strain Jc-1 was used. For infection of uPA-SCID mice with humanized liver HCV WT isolates from sera of an infected patient (gt1b, strain GLT1^38^) or of strain mH77c (gt1a^39^) was used. For HAV, genotype IA WT strains were isolated from patient stool samples. Huh7.5, HepG2 and HepaRG cells were infected with Hepatitis A virus strain HM175/18f, genotype IA. For HAV infection of humanized mice, virus of GLT1 serum was obtained from a male patient under immunosuppression with tacrolimus, mycophenolic acid and methylprednisolone because of status post re-liver-transplantation. The local ethics committee approved data collection and analysis (ethics vote: S-677/2020, Heidelberg University, ethics committee of the medical faculty). HAV WT was obtained from stool samples of two anonymous donors, through allowance of Apulia Region, Italy (DGR n. 565 from 2/04/2014).

### PRR Stimulation

For TLR3 exclusive stimulation, poly (I:C) High Molecular Weight (HMW; Invivogen, San Diego, California) was added to the supernatant of cells at 10 or 50 μg/ml. Six hours after stimulation, cells were harvested for RNA extraction and RT-qPCR. For stimulation of cytosolic PRRs (RIG-I, MDA5) and TLR3 as well, poly (I:C) was transfected into cells using Lipofectamine2000 (Life Technologies, Karlsruhe, Germany). For transfection of 1 well in a 24-well format, different dilutions from 0.001 to 1 μg poly (I:C) were incubated at room temperature with 0.1 to 0.5 μl Lipofectamine2000 reagent in 100 μl OptiMEM for 5 minutes, and then added to the cells. RNA was also isolated from these cells 6 hours after transfection.

### Statistics

Independent biological replicates are denoted with n-numbers. To test for significance, 2-tailed -paired or unpaired – *t-*test, or Welch’s test, were performed using GraphPad Prism 5 software (GraphPad Software, La Jolla, CA, USA). *P < .05; **P < .01; ***P < .001.

Independent biological replicates are denoted with n-numbers.

## RESULTS

### Upregulation of ISGs and chemokines upon HAV and HCV infection of SCID Alb/uPA mice

Little is known so far on the contribution of innate immunity to the mechanisms of clearance versus persistence of HAV and HCV infections. SCID Alb/uPA mice with humanized liver, which lack functional murine B and T lymphocytes^40^, are permissive for both viruses and therefore represent the only *in vivo* model allowing a side by side comparison of the cell intrinsic innate immune responses to HAV and HCV in absence of adaptive immunity. Using two different virus variants each, we found on average higher HAV RNA amounts in the liver compared to HCV (Supplemental Fig. 1A), despite comparable repopulation efficiency with human hepatocytes (Supplemental Fig. 1B). We then measured common ISGs activated by viral dsRNA downstream of the main PRRs in hepatocytes and playing key roles in establishing an antiviral response^11, 14^. Fluctuations among the different samples (Fig. 2A, 2C) were likely associated with different sacrification timings (Supplemental Table 1) and the individual condition of the mice upon tissue engraftment^40^. Yet, we detected in almost all cases substantial upregulation of ISGs and chemokines by Immunohistochemistry (IHC) (Fig. 1A, B, Supplemental Fig. 1 C, D, E) and by RT-qPCR (Fig. 1 C, D, Fig. 2 A – F) upon both HAV and HCV infection, as opposed to uninfected mice (Fig. 2G). Lower innate immune induction by HAV HM175/18f (Fig. 2A, 2C, 2F) correlated to its lower replication (Supplemental Fig. 1A).

**Figure 1.**
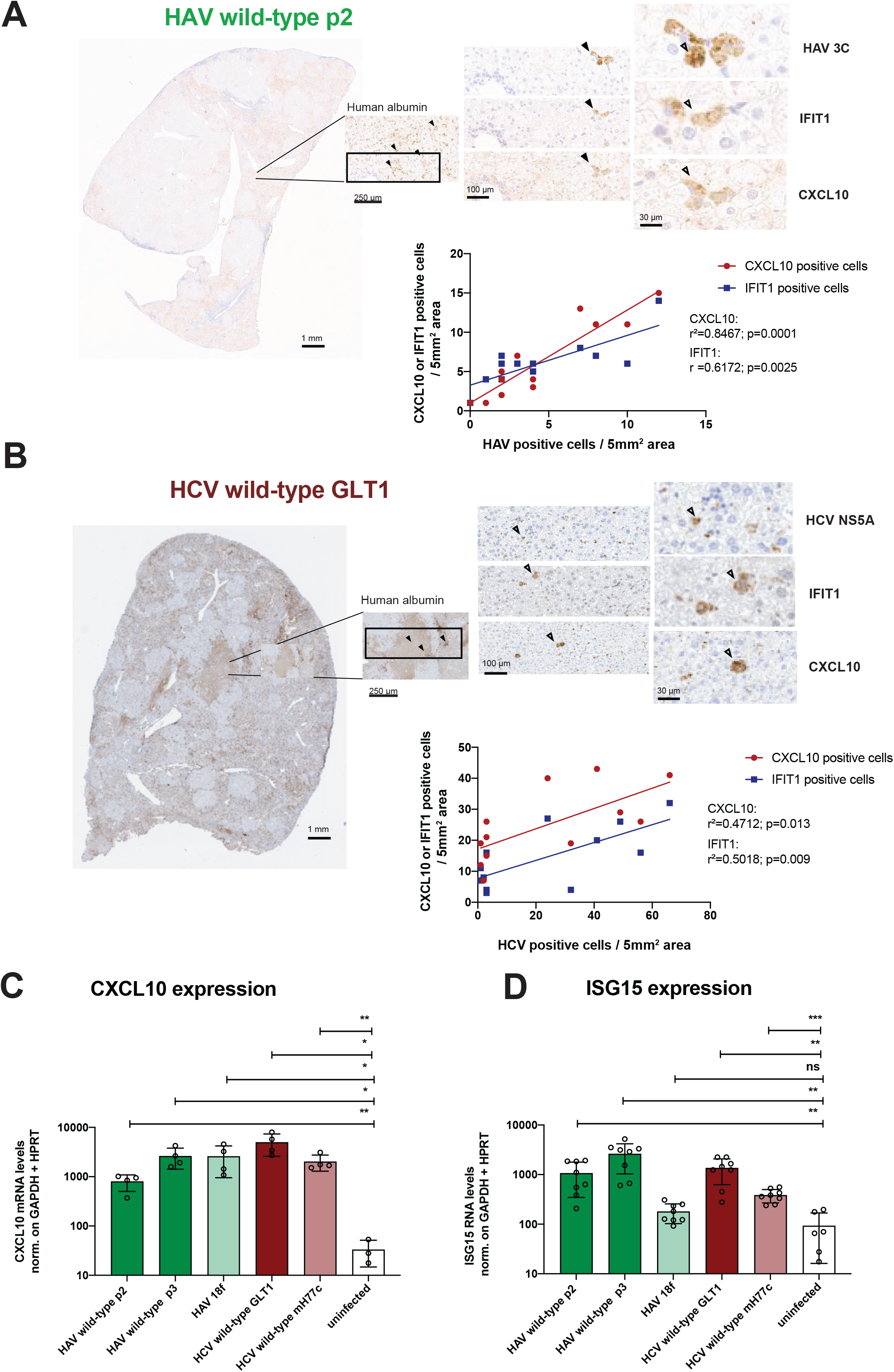
Upregulation of ISGs and chemokines upon HAV and HCV infection of SCID Alb/uPA mice. (A, B) Sections from the liver of SCID Alb/uPA humanized mice infected with wild-type HAV (A) or HCV (B) and subjected to IHC, using human albumin, HAV, HCV or ISG specific antibodies, as indicated. Shown are representative examples with abundant albumin signal (A, B, left and above panels). For each mouse, 6 albumin-rich view fields were quantified by hand for viral antigen positive cells as well as ISGs signals. Black arrowheads indicate a triple positive cell. Linear regression analysis was performed on CXCL10 or IFIT1 positive cells and HAV 3C or HCV NS5A positive cells, and statistical significance was assessed through Welsch’s unpaired *t-*test (A, B, lower right panels). (C, D) Total RNA was isolated from liver of uninfected mice or infected mice (n ≥ 3) and CXCL10 (C) or ISG15 (D) mRNA was quantified by RT-qPCR using human specific primers, and normalized to GAPDH and HPRT expression.

**Figure 2.**
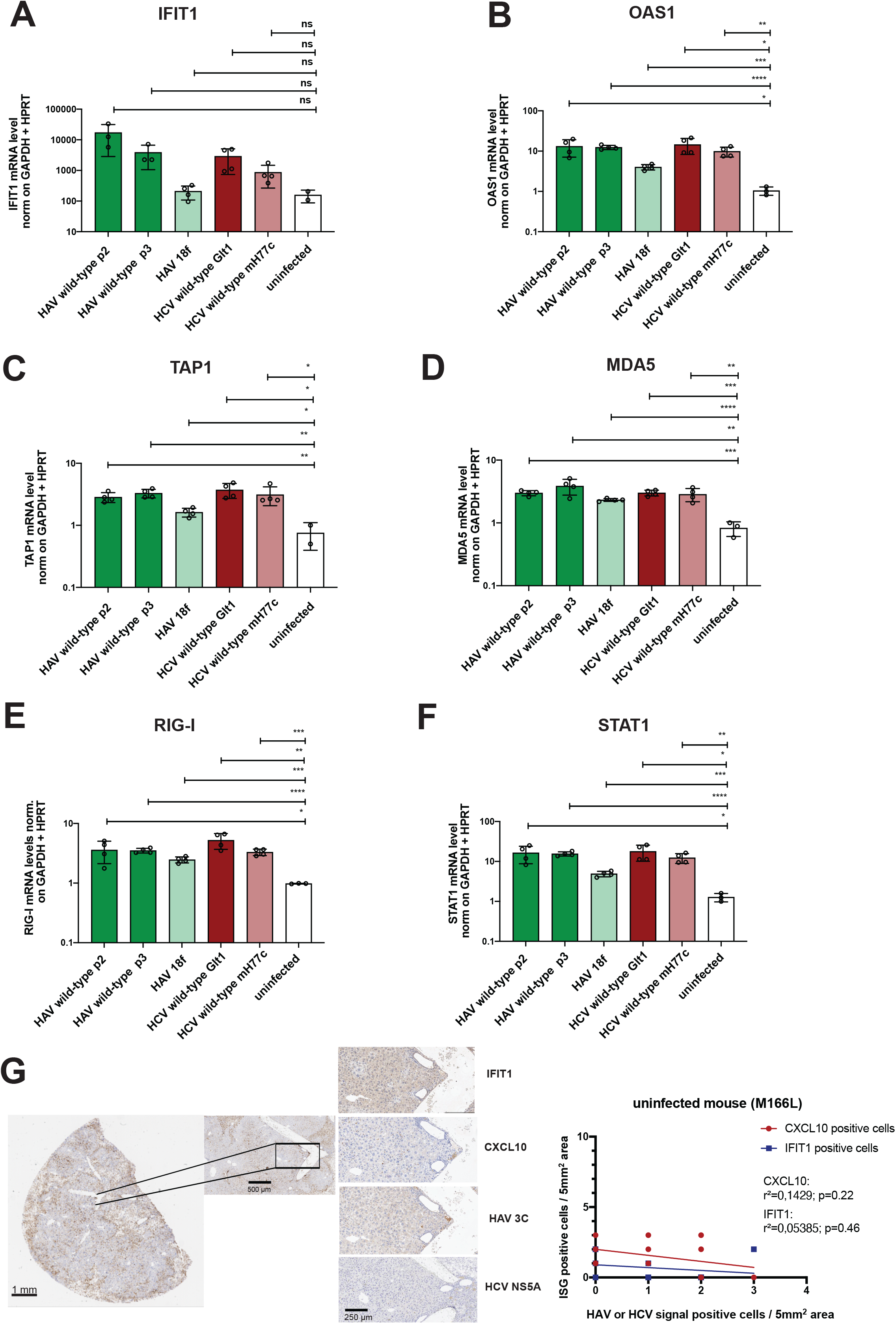
ISG induction in livers of HAV or HCV infected uPA-SCID mice with humanized liver compared to non-infected controls. (A – F) Total RNA was isolated from 20mg of liver of uninfected mice as well as from HAV or HCV infected mice (n ≥ 3) and mRNA expression levels of a set of ISGs and chemokines were quantified by a Taqman Gene Expression Array. Gene expression was normalized to GAPDH and HPRT expression. (G) Section from the liver of an uninfected uPA-SCID mouse with humanized liver subjected to IHC for HAV or HCV viral antigen signal and ISGs. Shown is an example of a repopulated area, characterized by abundant albumin signal, that was chosen for magnification (left and middle panel). 6 albumin-rich view fields were manually quantified for viral antigen positive cells as well as ISGs signals. Linear regression analysis was performed on CXCL10 or IFIT1 positive cells and HAV 3C or HCV NS5A positive cells, and statistical significance was assessed through Welsch’s unpaired *t-*test (right panel).

Overall, these data argued for a substantial cell intrinsic innate immune activation upon both Hepatitis A and Hepatitis C virus infections in the livers of humanized mice devoid of adaptive immunity.

### HAV does not trigger an innate immune response in Huh7 cells with reconstituted TLR3, MDA5 and RIG-I expression

Since previously only a very limited ISG response was described in the liver of three HAV infected chimpanzees^26^ we sought to determine the source of the strong innate immune induction detected by us in the HAV infected human liver chimeric mice. To allow direct comparison with HCV we used Huh7 cells, which are permissive for both viruses. All Huh7 variants lack detectable expression of TLR3 and MDA and contain only little or no functional RIG-I in case of subclone Huh7.5^10^. Therefore, reconstitution of PRRs relevant for dsRNA sensing is required to restore their signaling functions (Supplemental Fig. 2A, 2B). To validate induction of cell intrinsic innate immune responses by authentic viral replication intermediates we used Huh7 cells selected for the presence of HAV or HCV subgenomic replicons^36^ (Fig. 3A), in which abundant viral RNA was detectable (Fig. 3C) and transiently expressed TLR3, RIG-I or MDA5 (Fig. 3D). IFIT1 expression was induced in HCV replicon cells expressing TLR3, but not RIG-I and MDA5, in line with previous results^14^, whereas no induction by HAV was observed for any of the PRRs (Fig. 3D). In addition, we investigated the innate immune response in the context of infection with full-length genomes (Fig. 3B) on Huh7.5 cells stably expressing TLR3, RIG-I or MDA5 (Supplemental Fig. 2A, 2B). HAV and HCV replicated with similar efficiency in all cell lines (Fig. 3E). But while HCV trigged TLR3 robustly and RIG-I transiently, HAV again did not induce a detectable ISG response in any of the PRR-expressing cell lines (Fig. 3F and Supplemental Fig. 2C). Altogether, these data suggested that, despite robust replication in Huh7 cells, HAV did not induce an innate immune response or, alternatively, potently abrogated the innate signaling pathways, as reported^18, 19, 41^.

**Figure 3.**
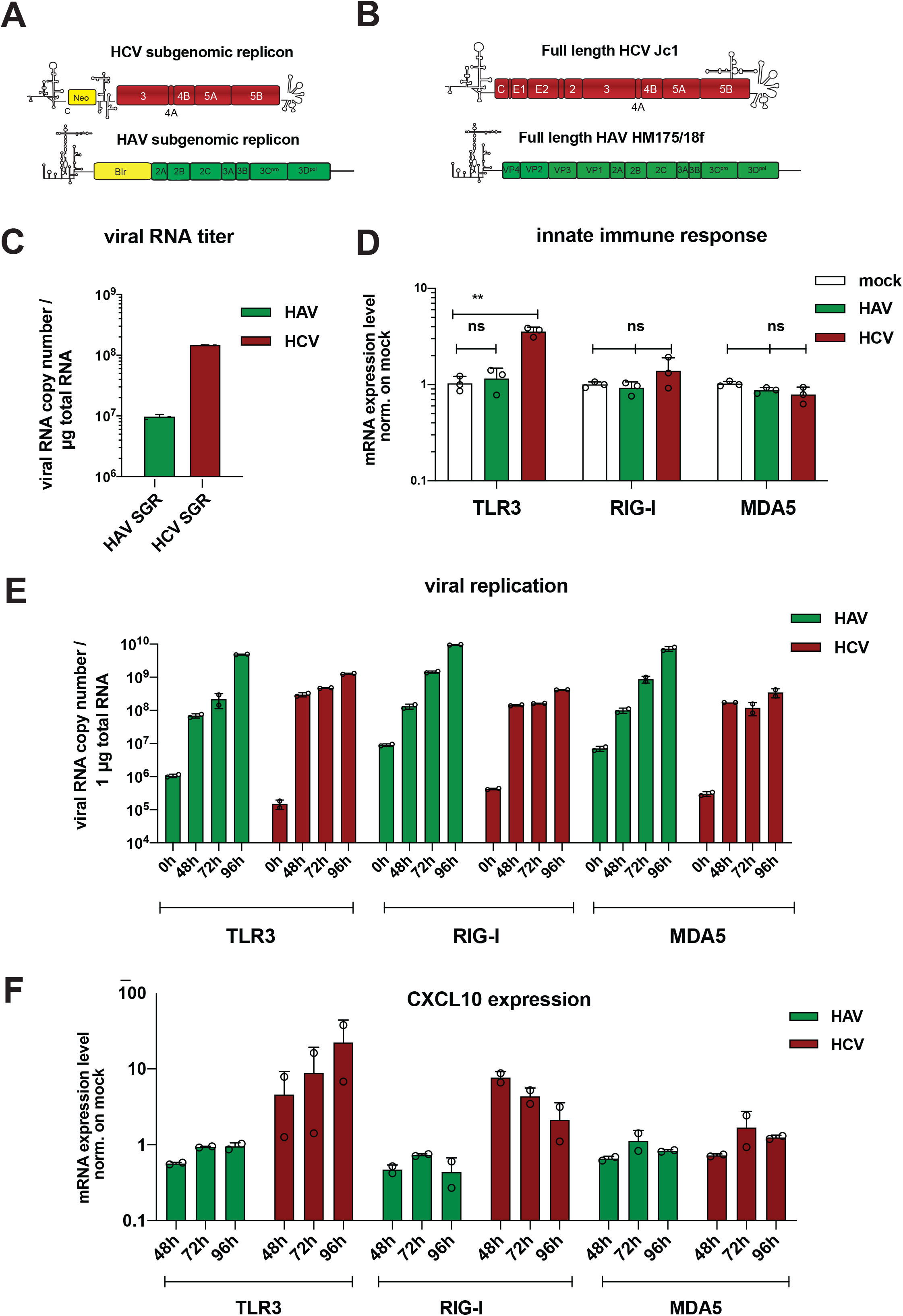
Innate immune response to HAV and HCV in Huh7 cells with reconstituted TLR3, MDA5 and RIG-I expression. (A) Schematic of the persistent, selectable subgenomic replicons used to select HCV / HAV stable cell lines (B) and the HCV / HAV genomes used for infection. Neo/Blr: genes conferring resistance to G418 or blasticidine, respectively. (C) HAV or HCV RNA was quantified in total RNA extracted from Huh7 cells harboring persistent subgenomic replicons by RT-qPCR. Viral copy numbers are expressed per μg RNA. (D) Different PRRs, as indicated, were transiently expressed in persistent replicon cell lines using lentiviral vectors. Twenty-four hours after transduction total RNA was isolated and IFIT1 mRNA was quantified by RT-qPCR and normalized to GAPDH expression. Data are shown as fold expression relative to untreated cells. Statistical significance was assessed by Welsch’s *t-*test. (E, F) Huh7.5 cells stably expressing either TLR3, RIG-I or MDA5, were infected with HAV (HM175/18f, green) or HCV (Jc1, red), respectively. At the indicated time points total RNA was isolated and CXCL10 mRNA (F), and viral RNA (E), were quantified as indicated. CXCL10 mRNA levels are normalized to GAPDH expression and shown as fold expression relative to uninfected cells. All values shown are mean values with SD from independent experiments (n = 2).

### The HAV 3C protease precursors cleave the adaptor proteins TRIF and MAVS only partially; HCV NS3-4A does not cleave TRIF, but fully degrades MAVS

Our work so far demonstrated that HAV induced innate immunity in human hepatocytes in vivo, but not in Huh7 cells. HAV replication in the latter was particularly robust, potentially resulting in a high expression of the viral protease 3C and the precursors 3CD and 3ABC, reported to cleave TLR3 and RLRs adaptors TRIF and MAVS, respectively^18, 19^. We first cloned TRIF, MAVS, the HAV and HCV proteases in individual expression vectors (Fig. 4A). Since we observed strong cell death upon overexpression of TRIF, we switched to a TRIF variant devoid of the RHIM domain^42^ which indeed rescued cell viability (Fig. 4B). Upon co-trasfection of TRIF and the HAV 3CD protease we detected the two expected cleavage products, concomitantly with a partial reduction of the full-length TRIF molecule upon 3CD wt, but not for a cleavage deficient protease mutant (Fig. 4C, black arrows). In contrast to literature, but in line with TLR3 activation by HCV replication^13, 14^, we could not detect any TRIF cleavage by HCV NS3-4A, neither for gt1b (Fig. 4D), nor for gt2a (Supplemental Fig. 3A). TRIF-cleavage was enhanced upon increasing amounts of 3CD expression, but not for NS3-4A (Supplemental Fig. 3B). To ensure expression of viral proteases in all cells, we also transfected TRIF in stable subgenomic replicon cells, detecting around 50% cleavage efficiency for HAV (Fig. 4E left) and almost none for HCV (Fig. 4E right). In infected cells, TRIF was again cleaved incompletely by HAV 3CD (Fig. 4F left) and not cleaved by HCV NS34A (Fig. 4F right).

**Figure 4.**
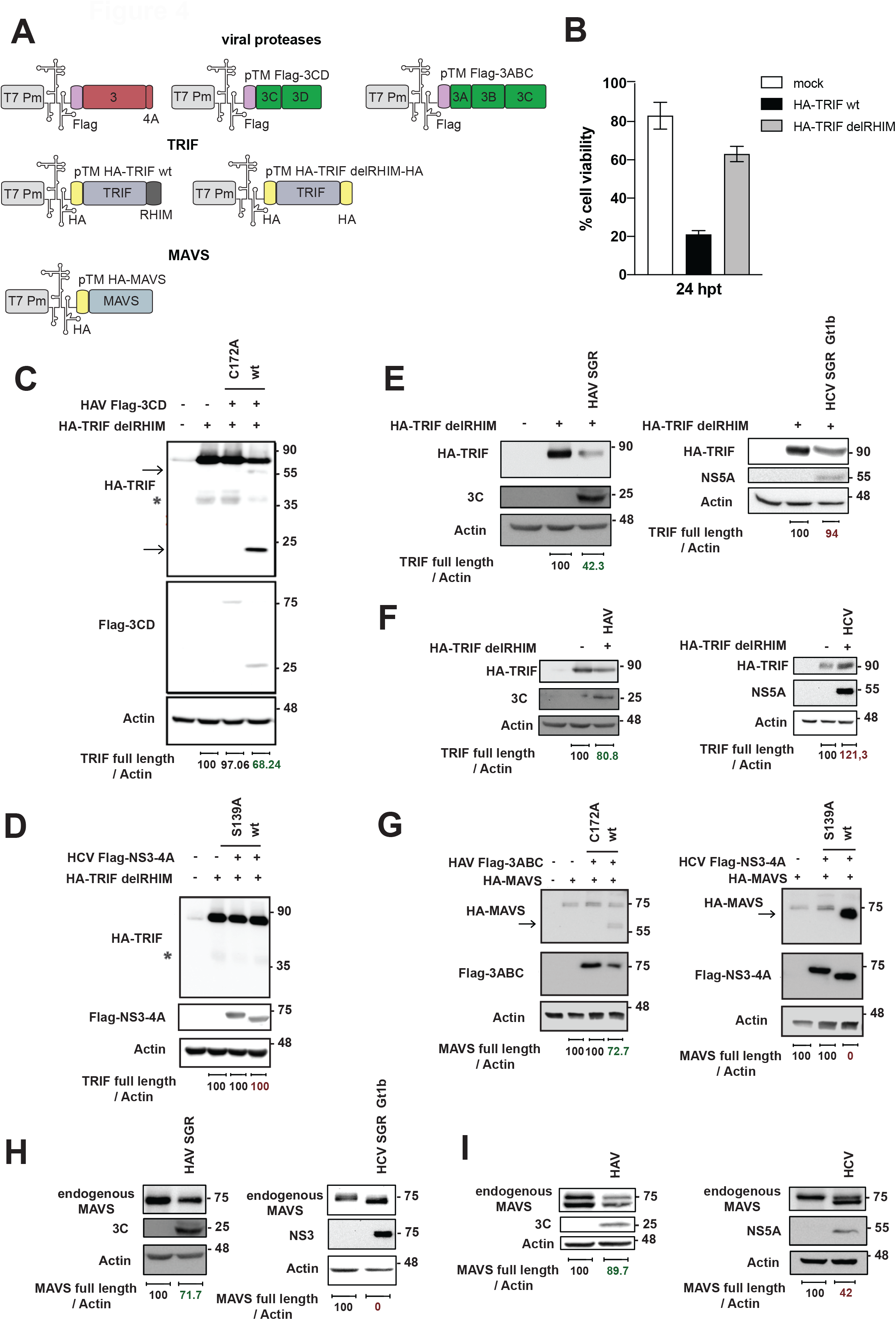
Analysis of TRIF and MAVS cleavage by HCV and HAV proteases. (A) Schematic of the expression vectors, encoding HAV/HCV viral proteases or the adaptor proteins TRIF / MAVS, used for transfection experiments. (B) Impact of the TRIF domain RHIM on cell viability. Huh7-Lunet T7 cells were transfected with a pTM vector encoding TRIF wt or TRIF delRHIM for 24 hours. WST-1 assay was used to determine cell viability. Shown are mean values and SD of triplicates from 1 experiment. (C, D, G) Huh7-Lunet T7 cells were co-transfected with plasmids encoding HA-TRIF delRHIM (C, D) and either FLAG-HAV 3CD (C) or FLAG HCV NS3-4A (D) or HA-MAVS (G) and either FLAG-HAV 3ABC (G, left panel) or FLAG-HCV NS3-4A (G, right panel) and harvested sixteen hours after transfection. Inactive mutants of HAV 3CD, C172A (C, G) and of HCV NS3-4A, S193A (D, G), were included. Arrows indicate specific cleavage products of viral proteases. Caspase-mediated TRIF cleavage products are indicated by an asterisk. (E) HAV (left) or HCV (right) persistent subgenomic replicon cells expressing T7 RNA polymerase were transfected with a plasmid encoding HA-TRIF under transcriptional control of a T7 promoter and harvested sixteen hours after transfection. Note that 3ABC and 3CD cleavage products in HAV replicon cell lines were not detectable, despite reduced amounts of full-length TRIF. (H) Endogenous expression of MAVS was detected in HAV (left) or HCV (right) persistent subgenomic replicon cells. (F, I) Huh7-Lunet T7 CD81h cells were infected with HAV (HM176/18f, F left, I left) and HCV (Jc1, F right, I right), respectively. Three days after infection, the cells were transfected with a plasmid encoding HA-TRIF under transcriptional control of the T7 promoter for detection of TRIF cleavage (F), or directly harvested for detection of endogenous MAVS cleavage (I). Note that 3ABC and 3CD cleavage products were not detectable in infected cells despite reduced amounts of the full-length protein (C-I). Lysates containing 10 μg of protein were analyzed by immunoblotting, using specific antibodies against HA, FLAG, HAV 3C, HCV NS5A, MAVS and β-actin (Actin), as indicated. Sizes of protein markers are indicated on the right [kDa]. Band intensities of TRIF or MAVS were normalized to the respective β-actin band intensities and quantified using Fiji.

As observed for TRIF, when we analyzed the activity of HAV 3ABC on MAVS cleavage upon transfection, we detected similar levels of partial cleavage, with full length MAVS still being the most prominent species (Fig. 4G, left). On the contrary, HCV NS3-4A fully cleaved MAVS, in line with literature^15-17^ (Fig. 4G, right). Full cleavage was obtained also with gt2a NS3-4A and abrogated by a mutation at the cleavage site (Supplemental Fig. 4C). Indeed, minimal HCV NS3-4A concentrations already cleaved MAVS at high efficiency, in contrast to HAV 3ABC (Supplemental Fig. 4D).

Since endogenous MAVS expression was well detectable, as opposed to TRIF, we were able to analyze MAVS cleavage in replicon cells and upon infection without ectopic expression. MAVS was again partly cleaved by HAV and fully by HCV NS3-4A in the replicon cells (Fig. 4H). In HCV-and HAV-infected cells MAVS cleavage efficiency was reduced for both viruses, likely due to the presence of uninfected cells (Fig. 4I). Therefore, we aimed at validating MAVS cleavage in a microscopy-based approach. To this end, we used GFP with a nuclear translocation signal, fused to the C-terminal membrane anchor of MAVS (MAVS-GFP-NLS)^43^. Compared to previous studies, we extended the MAVS coding region to encode the canonical protease cleavage sites of both viruses, allowing the quantification of nuclear GFP as a measure of protease cleavage. Both proteases were able to induce nuclear GFP translocation upon infection, yet to a higher degree for HCV (Supplemental Fig. 4A, 4B). Instead, upon transfection of the proteases, MAVS translocation upon 3ABC cleavage seemed impaired (Supplemental Fig. 4C, 4D), despite similar expression levels and localization of HAV 3ABC and HCV NS3-4A (Supplemental Fig. 4E, 4F), hinting at the cleavage upon infection being the most physiological of the three approaches we used.

In summary, these data showed that the HAV protease 3CD and 3ABC cleave TRIF and MAVS incompletely, and corroborated that HCV NS3-4A fully degrades MAVS, but does not cleave TRIF.

### Partial TRIF and MAVS cleavage by HAV protease precursors does not result in efficient interference with innate immune pathways

Having detected different cleavage efficiency by the HAV and HCV proteases, we next aimed to study the impact of TRIF and MAVS cleavage on functional counteraction of innate immunity. Therefore, we sought to generate stable cell lines expressing either the HAV or HCV protease, wild-type or mutant, along with the individual dsRNA sensor. Due to cytotoxicity of HAV 3CD (Supplemental Fig. 5A), we needed to rely on transient expression (Supplemental Fig. 5B). Next, we stimulated the cells with increasing concentrations of p(I:C) and quantified IFIT1 mRNA levels as a measure of ISG induction. Huh7 TLR3 cells were able to mount a full response to p(I:C), regardless of the presence of any viral protease (Fig. 5A). RIG-I and MDA5 responses were in contrast fully abrogated by HCV NS3-4A wt, but functional upon HAV 3ABC expression (Fig. 5B, 5C), despite comparable protease expression levels (Supplemental Fig. 5C). To include possible contributions of other viral proteins to the interference activity reported for HAV towards TLR3, RIG-I and MDA5, we next assessed ISG induction upon p(I:C) stimulation in subgenomic replicon cell lines reconstituted with each PRR, confirming that HAV proteases did not abrogate ISG induction by any of the PRRs, while HCV NS3-4A was blocking the RLRs (Supplemental Fig. 6).

**Figure 5.**
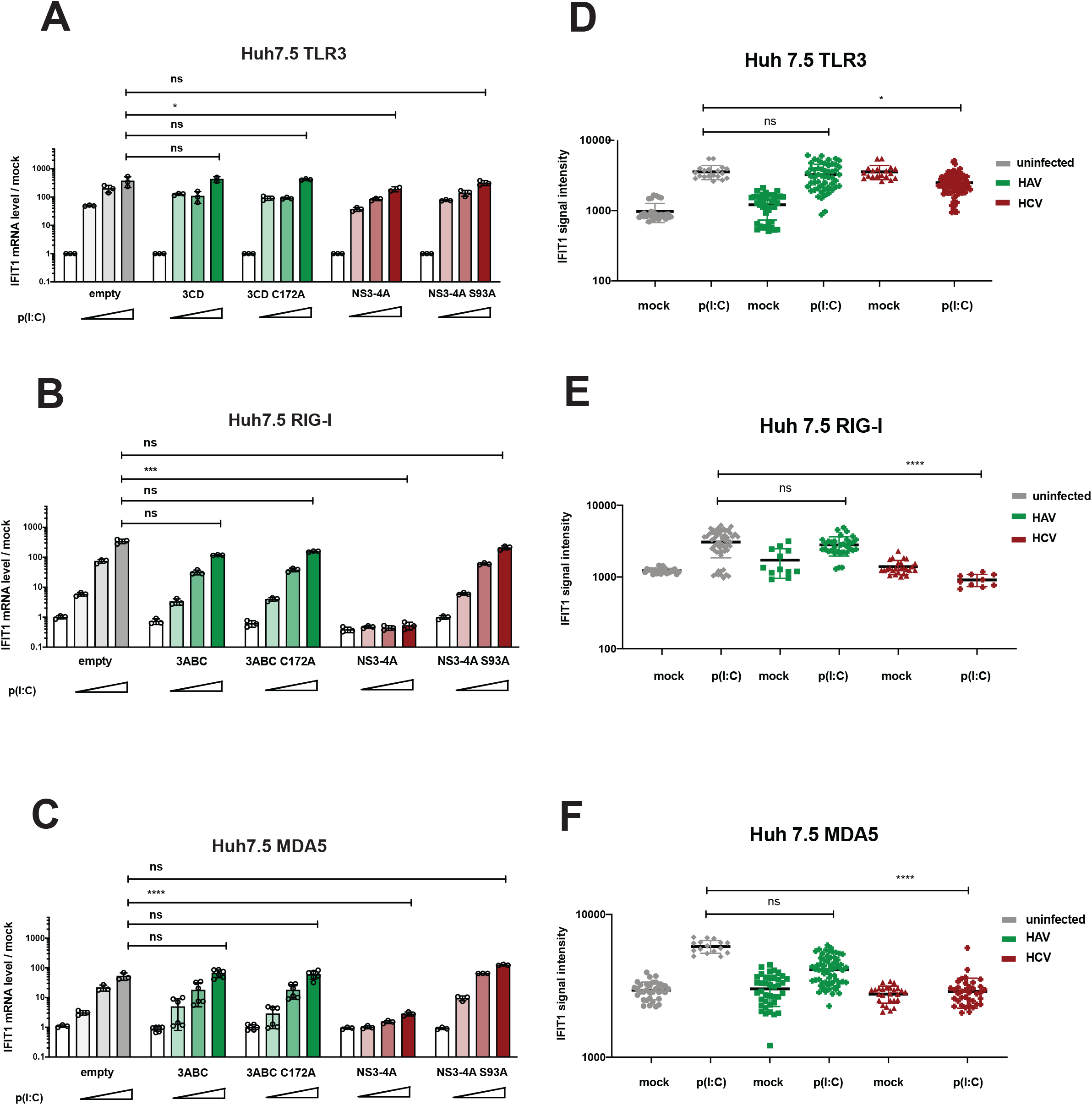
Impact of HAV and HCV protease mediated cleavage of TRIF and MAVS on cell intrinsic innate immune response. (A, B, C) Huh7.5 cells stably expressing either TLR3 (A), RIG-I (B) or MDA5 (C) were transduced with lentiviral vectors lacking inserts (empty) or encoding wt or mutant HAV proteases 3CD (A) or 3ABC (B, C) or HCV protease NS3-4A (A, B, C), as indicated. After selection and passaging, cells were transfected with increasing amounts of poly(I:C) (0, 0.01 μg/ml; 0.1 μg/ml; 1 μg/ml) and harvested six hours after stimulation to isolate total RNA and quantify IFIT1 mRNA by RT-qPCR. Data are normalized to GAPDH and shown as fold expression relative to uninfected cells. Mean values with SD from biological replicates (n = 3). Statistical significance was assessed by Welsch’s *t*-test. (D, E, F) Huh7.5 cells stably expressing either TLR3 (D), RIG-I (E) or MDA5 (F), were infected with HAV HM175/18f, green or HCV (Jc1, red), respectively. Three days after infection, cells were seeded onto coverslips and transfected with 0.5 μg poly(I:C) or mock treated (unstim). Six hours after stimulation, cells were fixed and IF staining was performed using viral antigen-(HAV Vp3 or HCV NS5A) and IFIT1-specific antibodies. Signal intensity for virus and innate immune response was quantified using Fiji and displayed in dot plots, where each dot represents a single cell. Statistical analysis was performed using Welsch’s *t*-test to compare the innate immune response to poly(I:C) in HAV or HCV infected cells compared to uninfected cells (grey).

We then asked whether this lack of counteraction could be influenced by the bulk measurement, potentially including cells with lower expression or no expression of the viral proteases. Thus, we established a single-cell analysis, infecting Huh7.5 cells reconstituted with individual PRRs and successively stimulating them with p(I:C), aiming at the detection of innate immune response in infected cells by IF, using IFIT1 specific antibodies (Supplemental Fig. 7A). Here, we did not detect a significant difference in IFIT1 level upon poly(I:C) stimulation between mock and HAV infected cells for all the PRRs (Fig. 5D, 5E, 5F and Supplemental Fig. 7B, 7C, 7D). In striking contrast, HCV infected cells completely abrogated RIG-I and MDA5 activation (Fig. 5E, 5F and Supplemental Fig. 7C, 7D). TLR3-reconstituted cells infected with HCV showed a higher IFIT1 even in absence of p(I:C), confirming the TLR3-specific sensing of HCV and the lack of strong counteraction by the virus (Fig. 5D and Supplemental Fig. 7B).

Altogether, we showed that TRIF and MAVS cleavages do not represent an efficient strategy of innate immune interference for HAV, as opposed to the strong HCV functional interference with the RLR pathways. Furthermore, we demonstrated that HCV did not counteract the TLR3 pathway through NS3-4A.

### HAV triggers innate immunity in cell culture models with an intact MDA5 pathway, essentially requiring LGP2

Having observed upregulation of ISGs in the HAV infected humanized mice, but not in Huh7 with reconstituted dsRNA receptors, we assumed that Huh7 cells might be defective for HAV sensing. LGP2 was shown to be an essential co-factor of MDA5 in mounting an IFN response^44, 45^ and MDA5 was reported to be the main sensor for HAV^11^. Thus, we assessed LGP2 mRNA expression in liver-based cell lines and indeed found that Huh7.5 cells lacked LGP2 expression (Fig. 6A). However, stable reconstitution of LGP2 expression in Huh7.5 MDA5 cells (Supplemental Fig. 8A) indeed increased ISG induction upon HAV replication (Fig. Supplemental Fig. 8B, C), but also induced higher baseline ISG expression in absence of infection, rendering these data difficult to interpret (Supplemental Fig. 8C).

**Figure 6.**
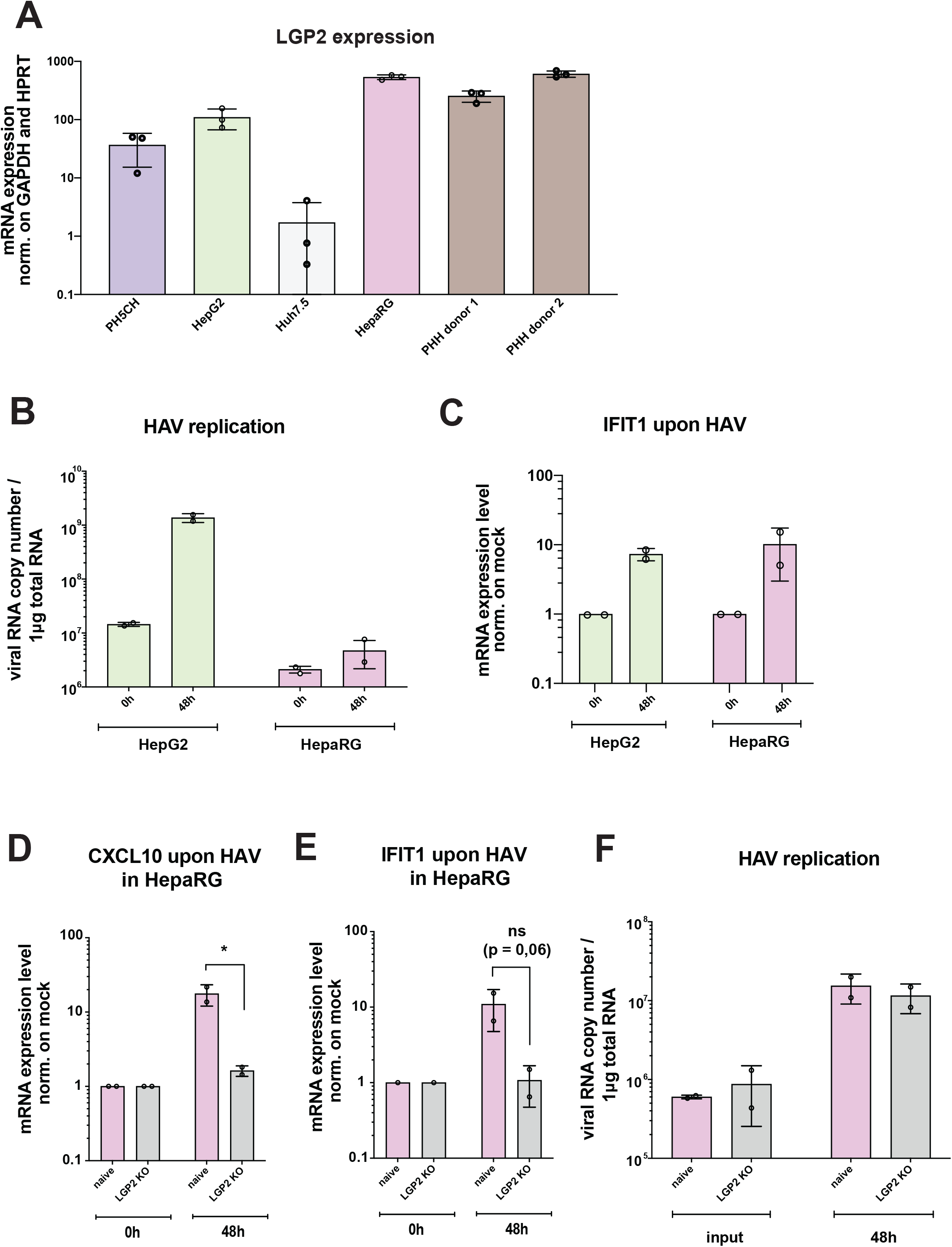
The sensing of HAV in hepatocytes requires LGP2 expression. (A) Total RNA was isolated from four different hepatocyte cell lines (PH5CH, HepG2, Huh7.5, and HepaRG) and Primary Human Hepatocytes (PHH) derived from two different donors. LGP2 mRNA levels were quantified by RT qPCR and normalized to GAPDH and HPRT expression. (B) HepG2 and HepaRG cells were infected with HAV HM175/18f. At the indicated time points total RNA was isolated and IFIT1 mRNA (C), and viral RNA (B), were quantified. IFIT1 mRNA levels are shown normalized to GAPDH expression relative to timepoint 0. (D-F) HepaRG cell pools with knockout for LGP2, or mock, were infected with HAV HM175/18f. CXCL10 (D) and IFIT1 (E) mRNA were quantified at the indicated time points, as well as viral RNA (F). IFIT1 and CXCL10 mRNA expression levels were normalized to GAPDH expression relative to infected cells at timepoint 0. All values shown are mean values with SD from biological replicates (n = 2).

Hence, considering alternative cell culture models, we chose HepG2 and HepaRG because of their high LGP2 expression (Fig. 6A). HepG2 seemed to be more permissive compared to HepaRG (Fig. 6B). Still, both cell lines showed a clear upregulation of IFIT1 upon HAV infection (Fig. 6C). Neither HepaRG nor HepG2 were permissive for HCV in our hands, the latter not even after restoration of miR122 and CD81 expression^46^, therefore we could not compare these data to HCV infection.

As HepG2 cells expressed TLR3 at undetectable levels, we restored this signaling pathway by ectopic expression (Supplemental Fig. 8D, 8E), but found similar degrees of IFIT1 induction upon HAV replication in presence or absence of TLR3, suggesting its limited role in HAV sensing (Supplemental Fig. 8F, 8G). Next, we used the same HepG2 TLR3 cells to generate knock-out (KO) pools of RIG-I or MDA5 (Supplemental Fig. 9A). Here, we showed that upon robust HAV replication (Supplemental Fig. 9B, C) the only PRR involved in detecting HAV was MDA5 (Supplemental Fig. 9D), whereas lack of RIG-I expression did not impact on IFIT1 induction (Supplemental Fig. 9E).

To further examine the role of LGP2 in HAV sensing, we infected LGP2 KO HepaRG cell pools^37^ with HAV, and found a significant reduction of CXCL10 (Fig. 6D) and IFIT1 (Fig. 6E), while HAV viral replication did not differ among the two cell lines (Fig. 6F).

In conclusion, our results indicate that HAV triggers an innate immune response in human hepatocytes *in vivo* and *in vitro*, in permissive models with intact signaling pathways. HAV replication was sensed by MDA5 and essentially required LGP2 expression. The moderate proteolytic activity observed by HAV towards MAVS obviously did not abolish sensing in HepG2 and HepaRG cells. We further excluded sensing of HAV by TLR3, questioning the functional significance of partial TRIF cleavage.

## DISCUSSION

This study provides the first side-by-side analysis of innate immune induction and counteraction by HAV and HCV in hepatocytes *in vivo* and in cell culture. So far, both viruses have been shown to cleave MAVS and TRIF by virus-encoded proteases, abrogating IFN responses. But while HCV in most *in vivo* studies induced a detectable ISG expression, HAV is regarded as not inducing IFN in infected hepatocytes, based on data from infected chimpanzees^26, 47^. However, our data demonstrate that both viruses induce a similar level of ISG expression in infected human hepatocytes *in vivo*. Our comprehensive cell culture studies suggest that HCV mainly triggers TLR3, due to the lack of TRIF cleavage, in contrast to efficient control of RLR signaling by cleavage of MAVS. We further show that HAV is exclusively sensed by MDA5 with support by LGP2, triggering an ISG response -due to only partial cleavage of MAVS – that is insufficiently blocked.

The innate immune response in the context of HAV is controversially discussed in literature. Some studies indicate that HAV barely induces innate immune activation in infected cells, due to the efficient disruption of PRR sensing pathways upon proteolytic cleavage of MAVS and TRIF by HAV proteases 3ABC and 3CD^18, 19^. This seemed plausible due to the limited ISG induction observed in the livers of HAV infected chimpanzees^47^. In contrast, a strong innate immune response upon HAV infection was described in PHH and HepG2 cells^11^. In addition, knockout of MAVS in mice is sufficient to allow HAV productive infection, suggesting that induction of innate immunity in hepatocytes is a clear restriction factor in this model^31^. While this was initially attributed to the incapability of HAV proteases to cleave murine MAVS, a very recent study showed that humanization of MAVS by restoration of the protease cleavage site in this model was not sufficient to allow HAV infection^32^. Since most of the discrepancies might be explained by the use of different models, we used a comprehensive set of approaches, including expression of the HAV proteases, persistent replicons and HAV infection in hepatic cells. All data agreed in detecting only a partial cleavage of MAVS and TRIF, not sufficient to counteract induction by p(I:C). Indeed, HAV replication induced ISGs in HepG2 and HepaRG, but not in Huh7 cells, a common model used in HAV research. This was attributed to the absence of LGP2, which we found essential for sensing of HAV by MDA5, underpinning the importance of choosing appropriate models. We excluded a role of TLR3 in HAV sensing, therefore the biological significance of partial TRIF cleavage remains to be determined. Importantly, HAV infection of uPA-SCID mice with humanized liver clearly showed ISG and chemokine production in infected human hepatocytes *in vivo*, to a similar extent as for HCV, in absence of adaptive immune responses. In chimpanzees, ISG induction was also observed but interpreted as induced by cytokine secretion of T-cells^26^. In fact, cytokine expression upon HAV infection was quite variable among the three chimpanzees analyzed, as it was for the HCV infected animals, suggesting that overlapping innate and adaptive immune responses might have impeded a thorough interpretation and masked the cell intrinsic induction by HAV replication in hepatocytes. Our data further demonstrate that neither cleavage of MAVS, nor other mechanisms counteracting IFN induction^18, 19, 22, 25, 41^ are capable of completely blocking HAV sensing, since HAV induces a clearly detectable ISG response in hepatocytes. However, since elimination of MAVS is a prerequisite to allow HAV infection of WT-mice^31^, we cannot exclude that HAV cleavage of MAVS or other counteraction mechanisms might attenuate the innate immune response, but to a lower extent than previously described.

For HCV, efficiency of MAVS cleavage by NS3-4A has been shown by many studies^15-17^, as well as the complete block of RLR-mediated ISG induction^16, 17^. Therefore, it served as an excellent control for the stringency of our models. However, there is controversy regarding the TLR3/TRIF pathway. While efficient TRIF cleavage has been reported, mainly based on *in vitro* translation models^20^, other studies indicated that TLR3 is still activated upon HCV infection^13, 14^ and could not reproduce TRIF-cleavage^21^. Studies on TRIF are generally hampered by barely detectable expression levels and high cytotoxicity upon ectopic expression^42^. Therefore, we included a RHIM-domain deletion mutant^42^ which did not alter susceptibility to protease cleavage and allowed robust detection. However, again implementing ectopic expression of NS3-4A, subgenomic replicons and virus infection, we could not detect any cleavage of TRIF, but instead TLR3 activation by HCV replication. Therefore, our data strongly argue against a general impact of TRIF cleavage to innate immune induction by HCV. Other mechanism interfering with TLR3 responses upon HCV infection, like secretion of dsRNA^14^ or NS4B-mediated degradation of TRIF^48^ also do not have the capacity to block TLR3 induction, but might rather weaken it, given the robust, TLR3-specific ISG response induced by HCV infection in cell culture. Sensing of HCV by TLR3 therefore might be the primary candidate for the ISG induction observed by us and others in uPA-SCID mice with humanized liver^49, 50^, HCV infected chimpanzees^26, 47^ as well as in infected hepatocytes of chronic HCV patients^28, 51^. However, in chronic HCV infection a strong variability in the level of ISG induction was found and attributed to polymorphisms in the IFNL locus^52^. The ability to produce IFNλ4 was strongly associated with high ISG levels and lack of HCV clearance^53^, likely by attenuation of T-cell responses^54^. The substantial contribution of host genetics to infection outcome highlights the complex interplay between innate and adaptive immunity in determining persistence and clearance of HCV infections.

Altogether, our data suggest a similar level of innate immune induction in HCV and HAV infected hepatocytes, originating from very divergent sensing and counteraction schemes. Limited induction of TLR3 accompanied by complete control of the RLR pathway might support persistence in case of HCV. Here, a certain level of innate immune response appears to favor persistence, as indicated by the impact of IFNλ4^53^. For HAV, inefficient counteraction, with MDA5/LGP2-mediated induction of innate immunity, appears to have no major impact on viral replication in the acute phase, but might on the long run support clearance. Overall, adaptive immune responses are the key for clearance of both viral infections^35, 55, 56^. A comprehensive understanding of the determinants of persistence versus clearance of HCV and HAV infections will therefore require fully immunocompetent animal models, which are currently not available.

## ACKNOWLEDGMENTS

The authors thank Rahel Klein, Ulrike Herian and Lieven Verhoye for excellent technical assistance. We are grateful to Vibor Laketa and the Infectious Diseases Imaging Platform (IDIP) for microscopy support. We thank Yuri Kusov for the HAV subgenomic replicon, Stanley Lemon for a plasmid encoding HAV full-length genome and Verena Gauss-Müller for the HAV 3C antibody. Huh7.5 cells and the HCV NS5A antibody “9E10” were generous gifts of Charles M. Rice. We thank Takaji Wakita for the JFH-1 isolate. In addition, we thank Zhenfeng Zhang, Ann-Kathrin Mehnert, Philipp Klein, Christopher Neufeldt and Mirko Cortese for helpful discussions.

