## Supplemental Material for "Comparison of HAV and HCV infections *in vivo* and *in vitro* reveals distinct patterns of innate immune evasion and activation"

**Contents:**

**Supplemental Materials and Methods**
**Supplemental References**
**Supplemental Tables 1-2**
**Supplemental Figures 1-9**

#### SUPPLEMENTAL MATERIAL AND METHODS

##### Cell Culture

Huh7 and HepG2 cell lines were cultured in Dulbecco's Modified Eagle Medium (DMEM; Life Technologies, Darmstadt, Germany), supplemented with 10% fetal calf serum (FCS, Capricorn Scientific, Germany), non-essential amino acids (Life Technologies, Darmstadt, Germany), 100 U/ml penicillin and 100 ng/ml streptomycin (Life Technologies) and cultivated at 37°C and 5% $CO_2$. HepaRG-derived cells were cultured in William's E Medium (Life Technologies, Germany), supplemented with 10% fetal calf serum (GE Healthcare, Germany), 100 U/ml penicillin, 100 µg/ml streptomycin (Sigma Aldrich, USA), 2 mM L-glutamine (Life Technologies, Germany), 5 µg/ml insulin (Sigma Aldrich) and 50 µM hydrocortisone (Sigma Aldrich). For culture experiments longer than 3 days, DMSO was always added to the final concentration of 1.5 %. All stably transduced cells were kept under selection pressure by addition of 1mg/ml G418 (Geneticin, Life Technologies), 1µg/ml puromycin (Sigma-Aldrich, Steinheim, Germany) or 5µg/ml blasticidin (Sigma-Aldrich).

Primary human hepatocytes (PHHs) were isolated and cultured as described before[1]. Tissue donors gave written informed consent for the experimental use of their specimens. The protocol was approved by the ethics commission of Hannover Medical School (#252-2008) and covered with reference number # S-161/2007.

##### HAV and HCV infection of Huh7.5, HepG2, HepaRG cells

Huh7.5 cells were seeded 16 hours prior infection in 12 well plates at a density of 0.8 * $10^5$/ well or 1.2 *$10^5$ / well for microscopy or RNA extraction, respectively. HepG2 cell lines were seeded 16 hours prior infection in 12 well plates at a density of 3.5 * $10^5$ / well. One hour prior infection, complete medium was replaced by FCS-depleted medium. Cells were inoculated with HAV and HCV diluted in FCS-depleted medium at a multiplicity of infection (MOI) 4 (for HAV) and 1 (for HCV) unless otherwise stated. Different MOIs were chosen for both viruses to reach comparable replication levels, judged by positive strand RNA detection in total RNA of infected cells. Three hours after infection, cells were washed with phosphate-buffered saline (PBS) and complete medium was added.

**HAV purification from stool samples**

Stool samples were obtained from two patients ("2" and "3") infected with HAV genotype IA, 12 to 13 days after the onset of clinical symptoms, and sequenced through Sanger sequencing targeting the VP1-P2A junction. The samples, alongside with a non-infected control, were diluted to a 10% v/v solution with PBS. Upon sequential pipetting and vortexing the sample was roughly homogenized and then sonicated with a sonicator (Branson) in 3 steps of 1 minute pulsing at 160 W and 1 minute resting on ice. The material was then centrifuged at 1,500 x g for 10 mins. The supernatant was serially passed through 5µm filters (CHROMAFIL Xtra PES filter, 25 mm, luer lock #ref 729242, Macherey-Nagel, Germany), then centrifuged again at 15,000 g for 15 minutes and then at 15,000 g for 30 minutes. Further filtering onto 0.8µm, 0.45µm and 0.22µm filter units (Whatman® Puradisc 13 syringe filter) using ensured isolation of the viral particles.

**HAV and HCV infection of uPA-SCID mice with humanised liver**

Human-liver chimeric mice were generated by transplantation of approximately $10^6$ primary human hepatocytes (donor C342 from Corning; L191501 from Lonza, Switzerland) into homozygous $uPA^{+/+}$-SCID mice as previously described[2]. Human albumin quantification in mouse plasma was used to assess the level of liver humanization. Mice ($n \geq 2$ per group) were infected by intrasplenical or intraperitoneal injection of the respective viral inoculum (serum containing HCV isolate GLT1[3] , serum containing HCV isolate mH77c[4], HAV HM-175/18f, HAV wild-type isolated from stool), as indicated in Supplementary Table 1. Blood plasma or feces were collected every 1 to 2 weeks and the plasma HCV RNA or feces HAV RNA load was determined by RealStar HCV RT-qPCR (Altona Diagnostics, Germany) following total nucleic acid extraction (NucliSENS EasyMag, BioMérieux).

**Immunohistochemistry (IHC)**

Liver samples were fixed in 4% paraformaldehyde and paraffin-embedded (FFPE). Tissue was cut into 2 µm sections from FFPE and cryo-preserved tissues were prepared and stained with Hematoxylin/Eosin or IHC antibodies. Epitope retrieval was performed by using citrate buffer for 20 (HCV NS5A) or 30 minutes (all other antibodies). Primary antibodies directed against viral and human proteins were HAV 3C (1:1500; kindly received from Verena Gauss Müller,

Lübeck), IFIT1 (1:400; D01, Abnova), CXCL10 (1:100; JA-1082, Invitrogen), HCV NS5A (1:4000; 9E10).

**RNA Extraction from livers of uPA-SCID mice with humanised liver**

After collection of the livers, 100 mg of tissue samples were preserved frozen in 1.5 ml RNAlater solution (ThermoFisher Scientific). To extract RNA, 20 mg of tissue were grinded to powder using a Dounce tissue grinder set (Merck, Germany) on dry ice. Total RNA from uPA-SCID mice with humanised liver was isolated using the Bio&SELL RNA-Mini Kit (Nuremberg, Germany) from 20 mg of liver tissue stored at -80°C in RNAlater solution (ThermoFisher Scientific) according to the manufacturer's protocol.

**Immunofluorescence analysis and Microscopy**

For immunofluorescence (IF) analysis, $2*10^4$ cells were seeded on coverslips in 24-well plates. Forty-eight hours after seeding, cells were fixed with 4% paraformaldehyde in PBS for 15 minutes at room temperature, and permeabilized with 50 µg/mL digitonin (Sigma-Aldrich, Steinheim, Germany) in PBS for 15 minutes on ice. After washing 3 times with PBS, primary antibody incubation was performed at room temperature for 1 hour, diluted in 3% bovine serum albumin/PBS. Primary antibodies used were anti-Flag (1:400, Sigma Aldrich - F1804-2MG), anti-3C (1:1500, kindly provided by Verena Gauss-Müller), anti HAV Vp3 (C22H, 1:250, Thermo Fisher Scientific, MA USA), and anti-NS5A (9E10, generous gift from Charles Rice, 1:150). After washing, secondary antibody incubation was performed for 45 minutes in the dark at room temperature. Secondary antibodies used were anti-mouse IgG2a-AlexaFluor488 and anti-rabbit Ig-AlexaFluor568 (dilution 1:1000; all from Invitrogen, Carlsbad, CA). After washing, nuclei were stained with 250 ng/mL 4′,6-diamidino-2-phenylindole (DAPI) (Invitrogen) in PBS for 1 minute, washed again and mounted onto microscopy slides. MitoTracker Deep Red (M22426) (ThermoFischer Scientific) was used for mitochondrial staining. Microscopy was performed on a Nikon Ti-Eclipse epifluorescence microscope with a ×40 oil immersion objective, on at least 30 cells per condition. For signal intensity analysis, background was removed from all channels and intensity was calculated using a Measure plug-in macro in Fiji.

**Quantitative Real-Time PCR (RT-qPCR)**

Total RNA was isolated from cultured cells or supernatant using the NucleoSpin RNA Plus kit (Macherey-Nagel, Düren, Germany). RT-qPCR was performed as described before[1]. For gene expression analysis, complementary DNA (cDNA) was generated from RNA samples using High-Capacity cDNA Reverse Transcription Kit (ThermoFisher Scientific, Waltham, MA), and was used for qPCR analysis with the 2x iTaq Universal SYBR Green supermix (Bio-Rad, Munich, Germany). Reactions were performed on a CFX96 Touch Real-Time PCR Detection System (Bio-Rad) as follows: 95°C for 3 minutes, 95°C for 10 and 60°C for 30 seconds. Primers used were S_GAPDH (GAAGGTGAAGGTCGGAGTC), A_GAPDH (GAAGATGGTGATGGGATTTC); S_IFIT1 (GAAGCAGGCAATCACAGAAA), A_IFIT1 (TGAAACCGACCATAGTGGAA), S_CXCL10 (GGCATTCAAGGAGTACCTCTCTC), A_CXCL10 (TGGACAAAATTGGCTTGCAGGA). Glyceraldehyde-3-phosphate dehydrogenase (GAPDH) or Hypoxanthine-guanine phosphoribosyltransferase (HPRT) were used as internal reference genes, and relative gene expression was determined using the 2-ΔΔCT method by normalizing to untreated samples. In the case of time course experiments, all samples were normalized to the untreated sample at the relative timepoint.

For analysis of HCV and HAV genomic RNA, 1-step RT-qPCR was performed using qScript XLT One-Step RT-qPCR ToughMix (Quanta Biosciences, Gaithersburg, MD), according to the manufacturer's instructions. In brief, 15 µL of reaction mixture contained 7.5 µL 2x enzyme/buffer mix, 1 µM of each HCV-specific primer (TCTGCGGAACCGGTGAGT and GGGCATAGAGTGGGTTTATCCA), 0.27 µM HCV-specific probe (AAAGGACCCAGTCTTCCCGGCAATT), or HAV-IRES- specific primers (GGTAACAGCGGCGGATATTGG and AGTCAATCCACTCAATGCATCCA) with a HAV specific probe (TGTTAAGACAAAAACCAATTCAACGCCGGA); 3 µL template RNA and RNase-free water. To determine absolute RNA amounts, a serial dilution of an RNA standard ($10^1$ to $10^8$ HAV or HCV RNA copies per reaction) was processed in parallel. Reactions were performed using the following program: 50°C for 10 minutes, 95°C for 1 minute, and 40 cycles as follows: 95°C for 10 seconds, 60°C for 1 minute.

**Taqman Gene Expression Assay on chimeric liver mice RNA**

For each infected group (e.g. HAV pat2) of mice, two mice were selected. From each mouse, 2 liver tissue blocks (stored in RNA-later buffer) were cut. Total RNA was extracted from 20-

30 mg of liver tissue by RNeasy plus mini kit, according to manufacturing conditions. A tissue lyzer (Qiagen) was used for grinding. cDNA was generated from RNA using Maxima H Minus First Strand cDNA Synthesis kit. qPCR was done by Taqman Gene Expression Assays using pre-designed primer/probe pair (no array set-up). qPCR protocol was performed on a LightCycler 480 using following conditions: hold 50°C (2 min), hold 95°C (20 seconds), denature 95°C (3 seconds), anneal/extend 60°C (30 seconds) à 40 cycles, cooling afterwards. Chosen reference genes with GUSB and HPRT1 because of high stability.

**Plasmid Constructs**

Plasmids encoding HCV or HAV replicons were described before[5], as well as the HCV Jc1 plasmid used for HCV virus production[6]. The HAV 18f plasmid used for HAV virus production was a kind gift from Stanley Lemon.

Plasmids used in the overexpression experiments were designed based on pTM vectors,[7] described in[8] with the cloned genes under translational control of an EMCV IRES. For HAV 3CD, the 3CD wt CDS was amplified from the HAV genome (HM-175/18f) and N-terminal FLAG-tagged using forward primer #1 (see Supplementary Table 2) and reverse primer #2. The C172A mutant was produced from the same template by overlap extension PCR using primers #1 + #4 and primers #2 + #3. PCR fragments were first cloned into vectors pWPI-Neo/Bla using XmaI and SpeI restriction sites. For cloning into pTM-1-2 we used NcoI and SpeI restriction sites, after amplification of the FLAG-tagged 3CD wt and mutant with primers #14 and #5.. For HCV NS3/4A, pWPI-GUN encoding untagged NS3/4A (Con1) wt and catalytically inactive mutant S139A (described before in [9]) were used for cloning into pTM-1-2, amplifying the NS3/4A CDS with an N-terminal FLAG-tag using primers #6 and #7 and NcoI and SpeI restriction sites. A pTM vector encoding Flag-tagged NS3/4A gt2a was based on JFH1, GenBank accession number AB047639[10]. For TRIF-encoding plasmids, we amplified the human TRIF wt CDS from a cloned TRIF cDNA. An N-terminal HA-tag was added using forward primer #8 and reverse primer #9. The C372R and the Q554A mutants were produced by overlap extension PCR from the same template using either primers #8 + #10 and #11 + #9 or primers #8 + #13 and #12 + #9. The C372R Q554A double mutant was produced by overlap extension PCR from HA-TRIF Q554A cloned into pTM-1-2 using primers #8 + #10 and #11 + #9. For cloning into pTM-1-2 and pWPI-Neo, PCR products and vectors were digested with XmaI and SpeI. For deletion of the RHIM domain and addition of an extra

C-HA tag, pTM HA-TRIF TRIFΔRHIM was produced using primers #8 and #15. N-terminally HA tagged MAVS and MAVS mutant C508A were generated based on pTM-MAVS[9] using primers #16 + #17 and #16 + #18, respectively, and restriction enzymes SmaI and SpeI. To generate MAVS-GFP-NLS, a pTM plasmid was cloned with a nuclear translocation signal with GFP fused to the C-terminal membrane anchor of MAVS (MAVS-GFP-NLS), where the MAVS coding region was extended to encode the canonical protease cleavage sites of both viruses (Q428 - C508).

Plasmids generated on a pWPI backbone with genes were used for generation of lentiviral vector particles. pWPI specifically encoding TLR3 Puro, RIG-I Blr and MDA5-Gun have been described before[1]. pWPI 3ABC and 3CD were cloned with forward primers #19 and #1, respectively, and reverse primer #2. To obtain 3ABC mutants, overlap PCR was performed using primers #19 and #4 and #3 and #2. For the same mutant in 3CD, primers #1 and #4 and #3 and #2 were used. All PCR products were digested with SmaI and SpeI. pWPI NS3-4A Neo and pWPI NS3-4A S193A Neo were already described before[9].

Constructs LentiCRISPRv2_RIG-I_KO_Puro[11] and LentiCRISPRv2_MDA5_KO_Puro (described in [12 13]) were used to achieve specific knock-out pools.

Phusion Flash High-Fidelity PCR Master Mix (ThermoFisher Scientific) was used for all cloning PCR reactions. Restriction digests were done according to the instructions of the manufacturer (NEB). Gel purification, PCR cleanup, Mini- and Maxi plasmid preparations were done using Macherey-Nagel NucleoSpin Gel and PCR Clean-up, NucleoSpin Plasmid and NucleoBond PC500, respectively. DNA ligation was performed with T4 DNA Ligase (ThermoFisher Scientific) in the respective buffer for at least 30 min at room temperature. Chemically competent DH5α *E. coli* were used for transformation. Sequences of all newly generated plasmids were confirmed by Sanger-sequencing by Eurofins Genomics, Germany.

**In vitro Transcription and Virus Production**

*In vitro* transcription of HCV viral RNA was performed as described elsewhere[5]. For HCV virus production, in vitro transcribed RNA of the Jc1 variant was transfected into Huh7.5 cells, supernatant was harvested at 72 hours after transfection and passed through a 0.45-µm filter (Whatman plc, Maidstone, UK). Virus stocks were concentrated by ultrafiltration using a centrifugal filter device (Centricon Plus-70; Millipore, Bedford, MA) and stored in aliquots at −80°C. Virus titers were determined by $TCID_{50}$ assay using Huh7.5 cells as described before[3] .

For HAV virus production, HAV full-length genome RNA transcript was generated by *in vitro* transcription on a linearized plasmid encoding the HAV strain HM-175/18f (kind gift by Stanley Lemon). 14 µg of the plasmid were digested with SmaI in a total volume of 100 µl at 37°C for 2h, and then purified using NucleoSpin® Extraction II kit (Macherey-Nagel). The DNA was then eluted in 63 µl of water and in vitro transcription was performed on 5 µg DNA in a total volume of 100 µl, using 12.5 µl of 25 mM ribonucleoside triphosphate (rNTP) and 80 U T7 polymerase (Promega), as well as 100 U RNAsin® Ribonuclease Inhibitor (Promega) and 20 µl of 5X RRL buffer (400 mM Hepes/KOH pH 7.5; 60 mM MgCl2; 10 mM Spermidin; 200 mM dithiothreitol (DTT) for 2 h at 37°C and then additionally overnight adding 2 µl T7 polymerase. To remove the DNA template, 20 µl of RNAse-free DNase (1 U/µl, Promega) were added and incubated for 30 min at 37°C. Next, the RNA was extracted by chloroform-phenol as described before[5]. *In vitro* transcribed RNA (HM-175/18f) was transfected via electroporation into Huh7.5 cells as described before[5]. Briefly, $1.5x10^7$ Huh7.5 cells were resuspended in 1ml cytomix and 10 µg of in vitro transcribed RNA were used for transfection of 400µl of single cell suspension. Cells were seeded as described before[5]. Cell lysates and supernatant were harvested at 11 days after transfection and subjected to 3 freeze / thaw cycles and passed through a 0.45-µm filter. Viral titer was determined through $TCID_{50}$.

**Production of Lentiviral Vectors and Selection of Stable Cell Lines**

Selectable lentiviral vectors were generated and used for transient transduction or for generation of stable cell lines as described before[14]. Briefly, HEK293T cells were seeded in 10 cm cell culture dishes ($5*10^6$ cells /10 ml DMEM) 16 hours prior transfection and fed from around one hour before transfection with FCS-depleted DMEM. A DNA mixture was prepared with the plasmids pSPAX2-Gag-Pol (5.14 µg), pMD2-VSVG (1.71 µg) and a pWPI vector encoding the gene of interest (5.14 µg) in 400 µL of Opti-MEM. The polyethylenimine (PEI) mixture was purchased by Polysciences Inc (23966-2, PA, USA) and diluted to a concentration of 1mg/ml. A transfection mixture was prepared with 36 µL of PEI in 400 µL of Opti-MEM. The plasmid mixture was then thoroughly mixed with the PEI mixture in a total of 800 µL Opti-MEM and incubated at room temperature (RT) for 20 minutes, then added to HEK293T cells drop-wise. Media was changed to complete DMEM after 6 hours. 48 hours and 72 hours after transfection, the supernatant was collected and filtered through a 0.45 µm filter to remove cells and cell debris and then was either aliquoted and stored at -80°C or directly added to target cells. Newly generated vectors were HAV 3ABC wild-type and mutant C172A (Puro), HAV 3CD wild type and mutant C172A (Puro) and HCV NS3-4A wild type and mutant S193A (Puro) (see "Plasmid constructs").

**DNA transfection**

For all overexpression experiments through pTM plasmid transfections, TransIT-LT1 Reagent (Mirus Bio LLC, Madison, Wisconsin) was used according to the manufacturer's protocol (Mirus Bio LLC). Briefly, $1.5*10^5$ Huh7-T7 cells were seeded in a 6-well plate in a 2 ml/well volume. One hour prior to transfection, cells were fed with FCS-depleted medium and a mix was prepared with 2.5 µg of DNA combined with 7.5 µl of TransIT-LT1 Reagent in a total of 250 µl Opti-MEM. After mixing completely by pipetting gently, the mixture was incubated at RT for 20 minutes and was added dropwise to the cells, which were harvested 16 hours after transfection.

**Knock-out of RIG-I and MDA5 in HepG2 cells**

To achieve knock-out pools, lentiviral vectors encoding Cas9 and RIG-I and MDA5-specific sgRNAs (see "Plasmid constructs") were used to transduce the respective target cells, as described before[15]. Upon selection with 1µg puromycin/ml (Sigma-Aldrich, Steinheim, Germany) the surviving cell pools were phenotypically checked for knock-out efficiency by immunoblotting.

**Immunoblotting**

50 µg cell extracts were separated on 10 or 12% SDS-PAGE and subjected to immunoblotting. After transfer to polyvinylidene difluoride membranes and blocking at room temperature in PBS containing 0.1% Tween 20 (PBS/Tween) and 5% skim milk for 1 hour, the polyvinylidene difluoride blots were incubated with antibodies specific for MDA5 (1:1000; Cell Signaling, Danvers, MA), RIG-I (1:1000; Adipogen, San Diego, CA), TLR3 (1:1000; Abcam), actin (1:4000; Sigma-Aldrich), HA tag (1:1000; Abcam), Flag tag (1:2000; Sigma Aldrich - F1804-2MG); 3C (1:200, kindly received from Verena Gauss Müller), NS5A (9E10, 1:5000; generous gift from Charles Rice), NS3 (#49, made in house, 1:1000) at 4°C overnight. After washing in PBS/Tween, the membranes were incubated with anti-mouse (1:10,000)

and anti-rabbit (1:5000) horseradish peroxidase antibodies (Sigma-Aldrich) at room temperature for 1 hour. Proteins were detected by using the ECL Plus Western Blotting Detection System (Pierce, GE Healthcare, Little Chalfont, UK) according to the instructions of the manufacturer. Signal was recorded using the Advance ECL Chemocam Imager (Intas Science Imaging, Goettingen, Germany).

**Cell Viability Assay**

Cell proliferation/viability was assessed by WST-1 assay (Cell-proliferation reagent WST-1; Roche, Basel, Switzerland) as described by the manufacturers' protocols in 96-well plates. Background was determined by measuring absorbance of medium in empty wells incubated with the reagent, and was subtracted from absorbance in experimental wells. Values were normalized to control samples.

**SUPPLEMENTAL TABLES**

**Supplemental Table 1: Infection of uPA-SCID mice with humanised liver**

| **Mouse ID** | **Inoculum** | **Sacrification date** | **HAV in feces (copies/ml)** | **HAV in plasma (copies/ml)** | **HCV in plasma (IU/ml)** | **Viral load detection** |
|---|---|---|---|---|---|---|
| M162L 071119 | HCV GLT1; 50 µL IS | 4/06/2020 12 wpi | | | 1,37E+07 | 10 wpi |
| M143L 201119 | HAV patient 2; 25 µL IS | 4/06/2020 12 wpi | 1,96E+10 | 1,74E+08 | | 8 wpi |
| M131RL 241119 | HAV patient 2; 25 µL IS | 4/06/2020 12 wpi | 1,13E+10 | 5,82E+08 | | 8 wpi |
| M150 160120 | HAV patient 3; 25 µL IP | 4/06/2020 8 wpi | 4,41E+09 | 3,24E+08 | | 6 wpi |
| M69L 160419 | HAV (HM175/18f); 100 µL IP | 27/11/2019 20 wpi | 5,50E+06 | 1,25E+07 | | 19 wpi |
| M69RL 160419 | HAV (HM175/18f); 100µL IP | 27/11/2019 20 wpi | 6,61E+06 | 5,83E+06 | | 19 wpi |
| M166R 090720 | HCV GLT1; 50 µL; IS | 26.11.2020 8 wpi | | | 5,02E+06 | 8 wpi |
| M174LR 100720 | HCV GLT1; 50 µL; IS | 26.11.2020 8 wpi | | | 1,28E+06 | 8 wpi |
| M216L 100720 | HAV patient 3 (dilution 1/10); IP 100µL | 26.11.2020 8 wpi | 9,81E+09 | 1,83E+08 | | 8 wpi |
| M216R 100720 | HAV patient 3 (dilution 1/10); 100 µL IP | 26.11.2020 8 wpi | 8,46E+09 | 8,48E+07 | | 8 wpi |
| M222 100720 | HAV patient 3 (dilution 1/100); 100µL IP | 26.11.2020 8 wpi | 2,65E+09 | 1,41E+08 | | 8 wpi |

| | | | | | | |
|---|---|---|---|---|---|---|
| M222L 100720 | HAV patient 3 (dilution 1/100); 100 µL IP | 26.11.2020 8 wpi | 2,58E+09 | 2,85E+08 | | 8 wpi |
| M230LL 050720 | HAV patient 3 (dilution 1/1000); 100 µL IP | 26.11.2020 8 wpi | 4,88E+09 | 5,01E+08 | | 8 wpi |
| M162L 071119 | HCV GLT1; 50 µL IS | 4/06/2020 12 wpi | | | 1,37E+07 | 10 wpi |
| M226L 280520 | HCV mH77c, 10^6 IU/ml (stock), 150 µL IS | 03/03/21, 14 wpi | | | 2,19E+06 | 13 wpi |
| M226LL 280520 | HCV mH77c, 10^6 IU/ml (stock), 150 µL IS | 03/03/21, 14 wpi | | | 3,71E+05 | 13 wpi |
| M124 030120 | control (no infection) | 26/03/20 | | | | |
| M124L 030120 | control (no infection) | 26/03/20 | | | | |

* IS = intrasplenically
* IP = intraperitoneally
* IV = intravenously

**Supplemental Table 2: Oligonucleotides used for plasmid cloning**

| # | Primer | Sequence 5‘ –> 3‘ |
|---|---|---|
| 1 | XmaI-Flag-3C_for | gatccccggggatcatggactacaaagacgatgacgacaagtcaactttggaaatagcaggactg |
| 2 | SpeI-stop-3D rev | ggatccactagttcatgaaaggtca |
| 3 | 3C C172A for | gcgaaggtcttcctggaatggccggtggggccttggtttc |
| 4 | 3C C172A rev | gaaaccaaggccccaccggccattccaggaagaccttcgc |
| 5 | SpeI-stop-3D rev | ggatccactagttcatgaaaggtca |
| 6 | NcoI-NS34A for | gatcccatggactacaaagacgatgacgacaagatggcgcctattacggc |
| 7 | SpeI-stop-NS34A rev | gatcactagttcagcactcttccatctc |
| 8 | SmaI_HA_TRIF for | gatccccgggcaccatgtatccctatgacgtccccgactacgcggccatggcctgcacaggc |
| 9 | SpeI TRIF rev | gatcactagtcgcggtcattctgcctcctgcg |
| 10 | TRIF C372R for | cctcctcctcctcctccttcatctactcctcgttcagctcacctgacccc |
| 11 | TRIF C372R rev | ggggtcaggtgagctgaacgaggagtagatgaaggaggaggaggaggagg |
| 12 | TRIF Q554A for | cctgcgggaacagagcgcacacctggacggtgagc |
| 13 | TRIF Q554A rev | gctcaccgtccaggtgtgcgctctgttcccgcagg |
| 14 | 3C shuttle pTM for | gatcccatggactacaaagacgat |
| 15 | TRIF dRHIM_Spe C-HA rev | gatcactagttcaggccgcgtagtcggggacgtcatagggataggactgcgggaaggg |
| 16 | SmaI HA MAVS forward | gatccccgggcaccatgtatccctatgatgacgtccccgactacgcggccatgccgtttgctgaagacaag |
| 17 | MAVS wt 3’ reverse | gcccgtagtttactagttcatttcattc |
| 18 | MAVS C508R 3’ SpeI reverse | gcccgtagtttactag actagt catctactc |
| 19 | XmaI-Flag-3A_for | cccggggatcatggactacaaagacgatgacgacaag ggaatttcagatgataatgatagtgcag |

### SUPPLEMENTAL FIGURES

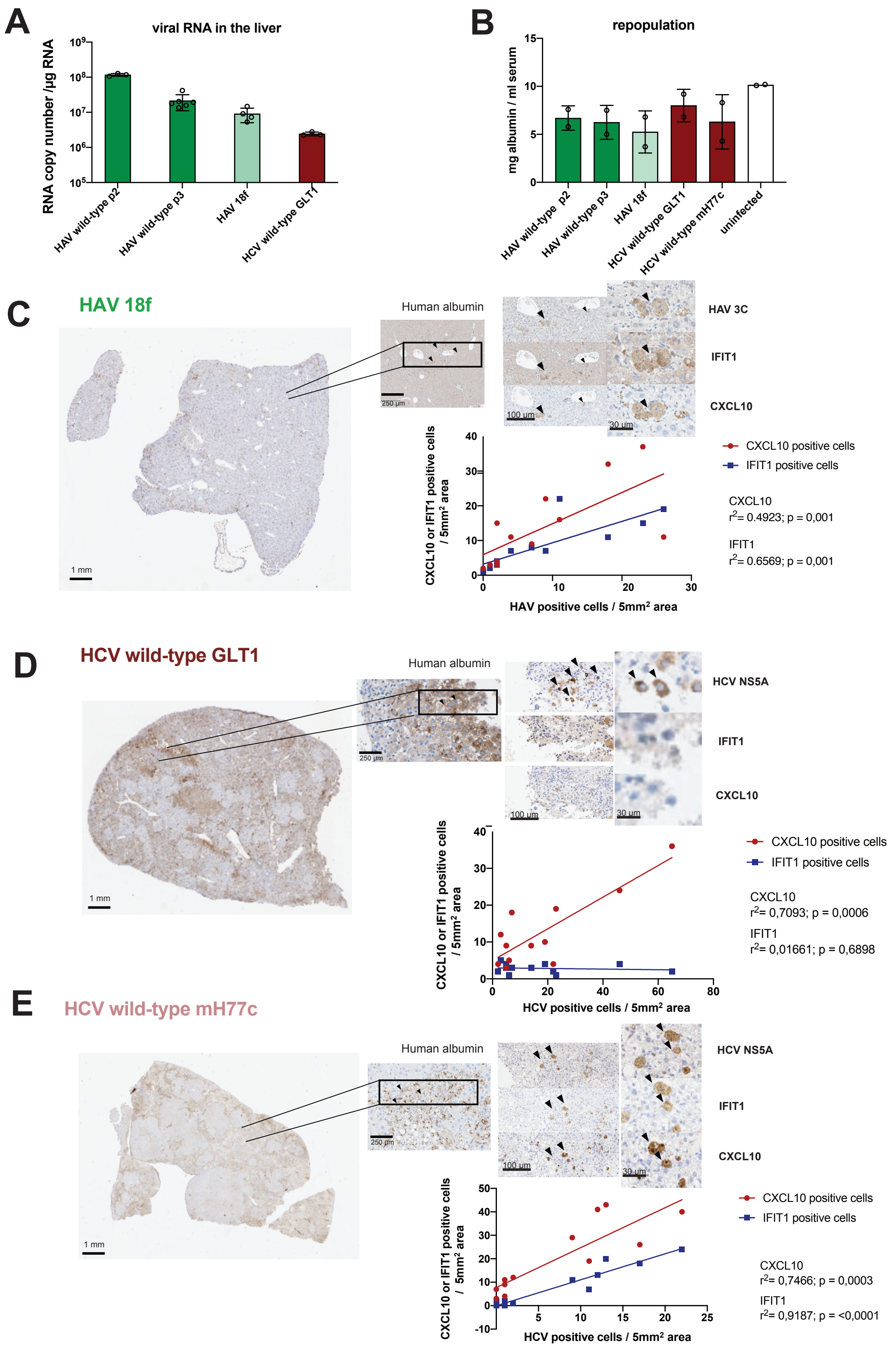

**Supplemental Figure 1. HAV and HCV infection uPA-SCID mice with humanised liver.**
(A) Viral replication in uPA-SCID mice with humanised liver. Total RNA was isolated from 20 mg of liver of HAV or HCV infected mice (n ≥ 3) and harvested at the time points indicated in Supplementary Table 2. Viral RNA was quantified by RT-qPCR using HAV and HCV specific primers. (B) Human tissue repopulation in uPA-SCID mice with humanised liver. Human albumin was measured in serum of HAV and HCV infected mice as well as in uninfected mice 6 to 8 weeks after engraftment.
(C, D, E) Consecutive sections from the liver of uPA-SCID mice with humanised liver infected with cell culture adapted HAV strain HM-175/18f (C), serum derived wild-type HCV Gt1b (isolate GLT1) (D) and serum derived wild-type HCV Gt1a (isolate mH77c) (E) were subjected to IHC for human albumin, HAV 3C or HCV NS5A and ISGs (IFIT1 or CXCL10), as indicated. Shown are examples of one repopulated area for each mouse, characterized by abundant human albumin signal, that was chosen for magnification (C, D, E, left and above panels). For each mouse, 6 albumin-rich view fields were quantified by manual counting for viral antigen positive cells as well as ISGs signals. Black arrowheads indicate a triple positive cell. Linear regression analysis was performed on CXCL10 or IFIT1 positive cells and HAV 3C or HCV NS5A positive cells, and statistical significance was assessed through Welsch's unpaired *t*-test (C, D, E, lower panel).

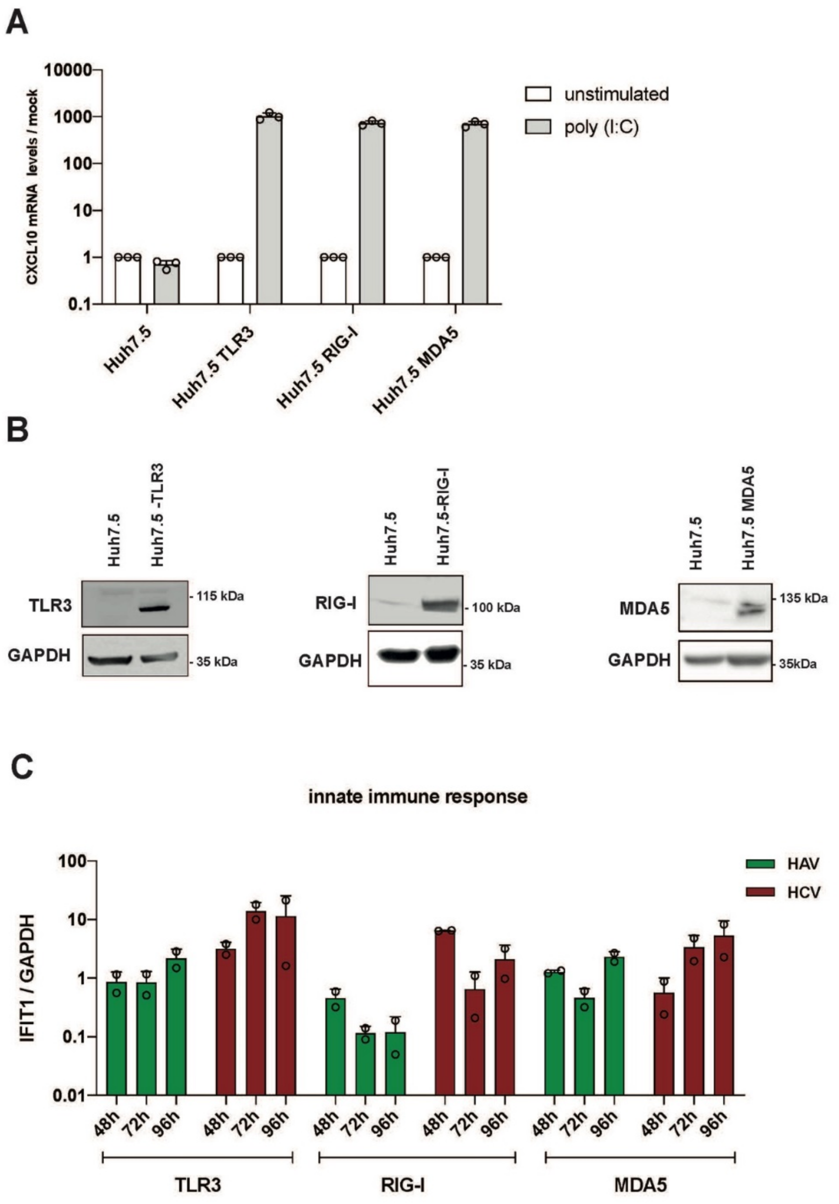


**Supplemental Figure 2. ISG induction in Huh7.5 cells reconstituted with PRRs and stimulated with p(I:C) or by infection with HAV or HCV.**

Huh7.5 cells were transduced with either TLR3, RIG-I or MDA5 using lentiviral vectors and selected and passaged for stable expression. (A) Huh7.5 cells expressing the indicated PRR were transfected with 0.5 µg/ml p(I:C). Six hours after stimulation, cells were harvested and RNA was isolated. Innate immune response was evaluated measuring CXCL10 mRNA levels via RT qPCR, shown normalized to GAPDH expression. Data are shown as fold expression relative to the unstimulated empty cells. Shown are mean values with SD from technical triplicates from 1 experiment (n=1). (B) Cell lysates from Huh7.5 cells, either empty or stably expressing the indicated PRR, were subjected to immunoblot analysis, using antibodies specific for TLR3, RIG-I or MDA5, as indicated. (C) Huh7.5 cells stably expressing the indicated PRR were infected with HAV (MOI 4, green bars) or HCV (MOI 1, red bars). Total RNA was extracted at the indicated time points post infection and analyzed for IFIT1 and GAPDH mRNA expression by RT-qPCR. Data are normalized to GAPDH expression and expressed as fold induction relative to uninfected cells. All values shown are mean values with SD from independent experiments (n = 2).

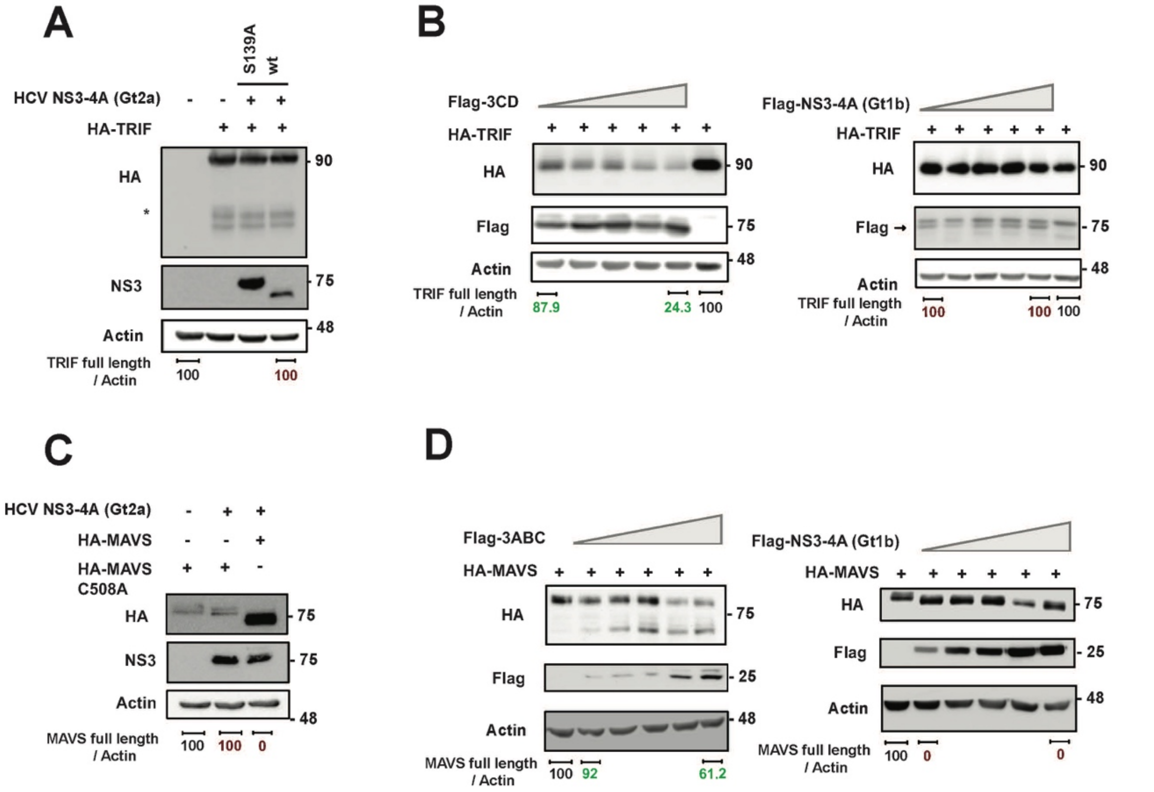


**Supplemental Figure 3. Cleavage of TRIF and MAVS by HAV and HCV proteases.**
Huh7-Lunet T7 cells were co-transfected with pTM vectors encoding HA-tagged innate immune adapter proteins MAVS or TRIF and Flag-tagged viral proteases. Sixteen hours after transfection, cell lysates were harvested and 10 µg of protein lysate was separated by SDS-PAGE and subjected to immunoblotting, using specific antibodies against HA-tag, Flag-tag or b-actin (Actin), as indicated. (A) 1µg of a plasmid encoding HA-TRIF was co-transfected increasing amounts of plasmids encoding either FLAG-HAV 3CD (left panel) or FLAG HCV NS3-4A (right panel), as indicated by the grey triangle (0.05 µg; 0.1 µg; 0.25 µg; 0.5 µg; 1.25 µg). (B) Plasmids encoding NS3-4A of HCV gt2a (JFH1), either wild-type or inactive mutant (S193A) and HA-TRIF were co-transfected. Bands associated with caspase-mediated TRIF cleavage are indicated by an asterisk. (C) 1µg of a plasmid encoding HA-MAVS was co-transfected increasing amounts of plasmids encoding either FLAG-HAV 3ABC (left panel) or FLAG HCV NS3-4A (right panel), as indicated by the grey triangle (0.05 µg; 0.1 µg; 0.25 µg; 0.5 µg; 1.25 µg). (D) Plasmids encoding HA-MAVS or a cleavage-resistant MAVS mutant (C508A) and HCV NS3-4A (gt2a, JFH1) were co-transfected. NS3-4 sequences are based on isolate Con1 (gt1b), unless otherwise indicated. Sizes of protein markers are indicated on the right [kDa]. Band intensities of TRIF or MAVS were normalized to the respective b-actin band intensities and quantified using Fiji.

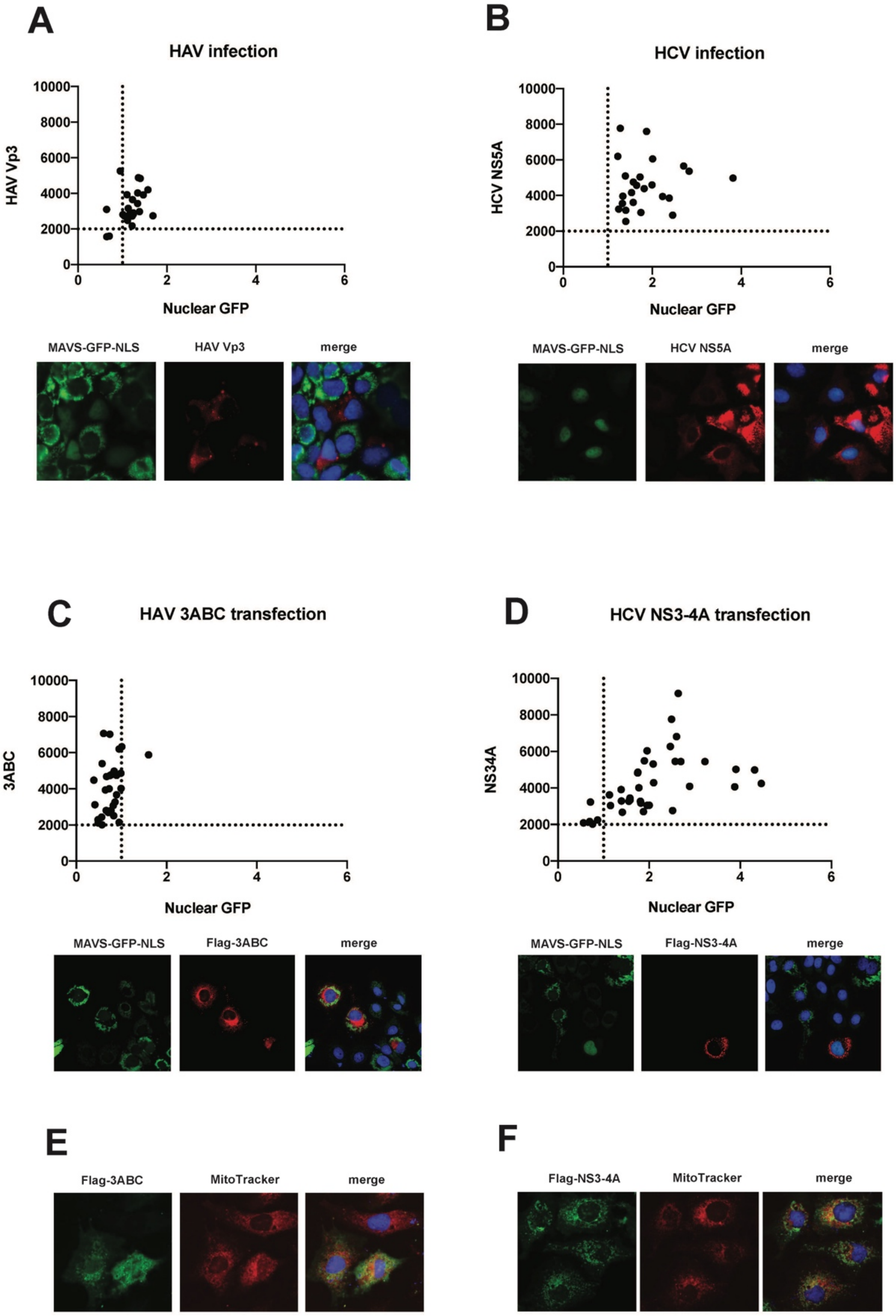
A
HAV infection
HAV Vp3
Nuclear GFP
MAVS-GFP-NLS
HAV Vp3
merge
B
HCV infection
HCV NS5A
Nuclear GFP
MAVS-GFP-NLS
HCV NS5A
merge
C
HAV 3ABC transfection
3ABC
Nuclear GFP
MAVS-GFP-NLS
Flag-3ABC
merge
D
HCV NS3-4A transfection
NS34A
Nuclear GFP
MAVS-GFP-NLS
Flag-NS3-4A
merge
E
Flag-3ABC
MitoTracker
merge
F
Flag-NS3-4A
MitoTracker
merge

**Supplemental Figure 4. Assessment of MAVS-cleavage by HAV and HCV proteases based on nuclear translocation of GFP.**

(A - D) Huh Lunet cells were transduced with lentiviral vectors encoding GFP tagged with a nuclear translocation signal (NLS) fused to the C-terminal membrane anchor of MAVS (MAVS-GFP-NLS), where the MAVS coding region was extended to encode the canonical protease cleavage sites of both viruses, allowing the quantifying of nuclear GFP as a measure of protease cleavage. Nuclear GFP signals were measured using Fiji in $n \geq 30$ cells. The ratio nuclear / cytoplasmic signal is expressed, along with the intensity of the signal from the viral antigen, as dot plot with each dot representing a single cell. (A, B) Cells stably expressing MAVS-GFP_NLS were seeded onto coverslips and infected with either HAV (A) (MOI 4) or HCV (MOI 1) (B). After three days, cells were fixed and stained with IF for viral antigens und with DAPI to stain nuclei. (C-D) Huh Lunet T7 cells, stably expressing a MAVS-GFP-NLS encoding the HAV and HCV cleavage sites, were transfected with 1µg of either FLAG-HAV 3ABC (C, E) or FLAG HCV NS3-4A (D, F). Eight hours after transfection, cells were fixed and stained for Flag-tag to assess viral antigen and DAPI. The ratio nuclear / cytoplasmic signal is expressed, along with the intensity of the signal from the Flag-tagged protease, as dot plot with each dot representing a single cell. (E, F) Huh Lunet T7 cells were transfected with 1µg of either FLAG-HAV 3ABC (E) or FLAG HCV NS3-4A (F). Eight hours after transfection, cells were fixed and stained for Flag-tag and mitochondria with MitoTracker Deep Red (M22426).

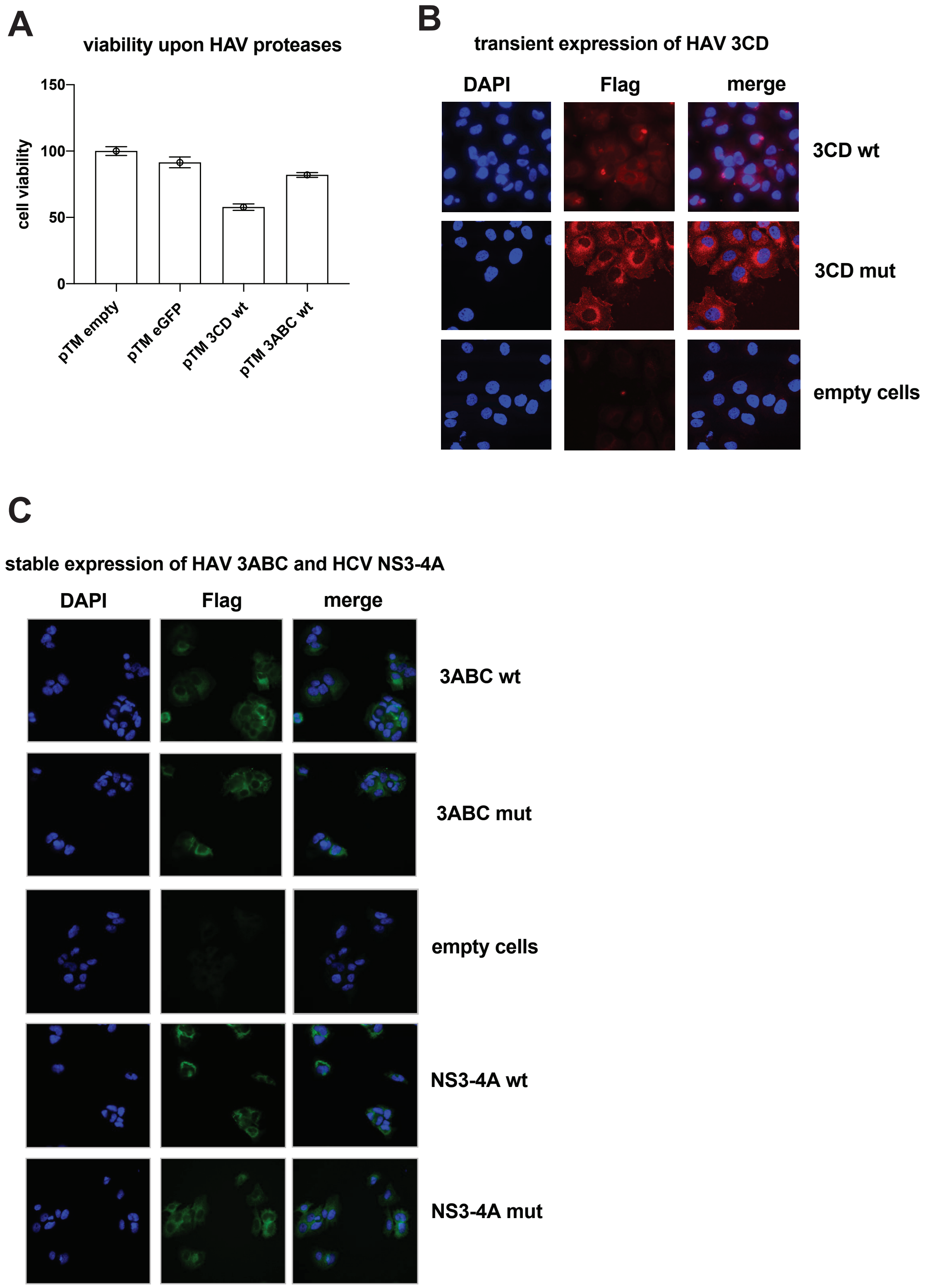


**Supplemental Figure 5. Cytoxicity of HAV proteases and detection of their expression upon transient or stable transduction with lentiviral vectors.**

(A) Huh Lunet T7 cells were transfected with pTM plasmids encoding GFP or HAV protease precursors 3CD wild-type and 3ABC wild-type. 24 hours after transfection cell viability was determined by WST-1 assay. Cell viability was normalized on cells transfected with an empty vector. Shown are mean values and SD of triplicates from 1 experiment. (B) HAV 3CD wild-type and mutant were transiently expressed in Huh7.5 cells using lentiviral vectors. 24 hours after transduction, cells were fixed and stained with a Flag-tag specific antibody and nuclei were stained with DAPI. (C) Stable expression of HAV 3ABC and HCV NS3-4A, both wild-type and inactive mutant, was achieved in Huh7.5 cells through transduction with lentiviral vectors. After selection and passaging, the cells were fixed and stained for Flag-tag and nuclei were stained with DAPI.

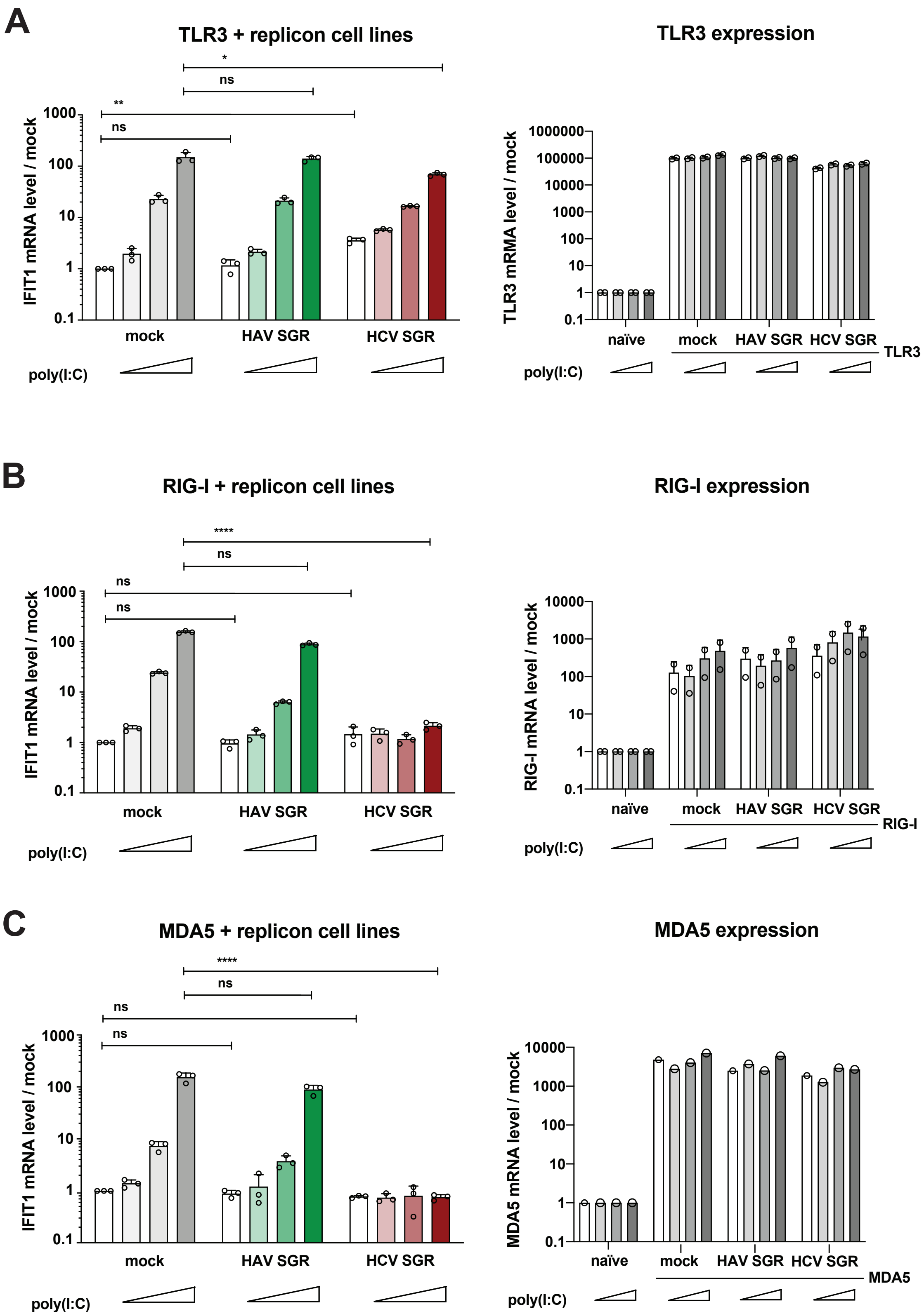


**Supplemental Figure 6. Interference with p(I:C) mediated innate immune induction in cell lines harboring persistent replicons.**

Lentiviral vectors were used to stably express TLR3 (A), RIG-I (B) or MDA5 (C) in Huh7 cells harbouring subgenomic replicons (SGR) of HAV or HCV, or in naïve Huh7 cells (mock). After transduction, cells were selected and subsequently stimulated by transfection of increasing amounts of p(I:C) (0.001 µg / ml ; 0.01 µg/ ml ; 0.1 µg /ml). Six hours after transfection, total RNA was isolated, then IFIT1 mRNA was quantified by RT-qPCR and normalised to GAPDH expression (A, B, C, left panels). Data are shown as fold expression relative to untreated cells. All values shown are mean values with SD from independent experiments (n = 3). Successful reconstitution of PRRs expression was confirmed in one representative experiment by RT-qPCR, and shown in A, B, C (right panels). Statistical significance was assessed by Welsch's *t*-test.

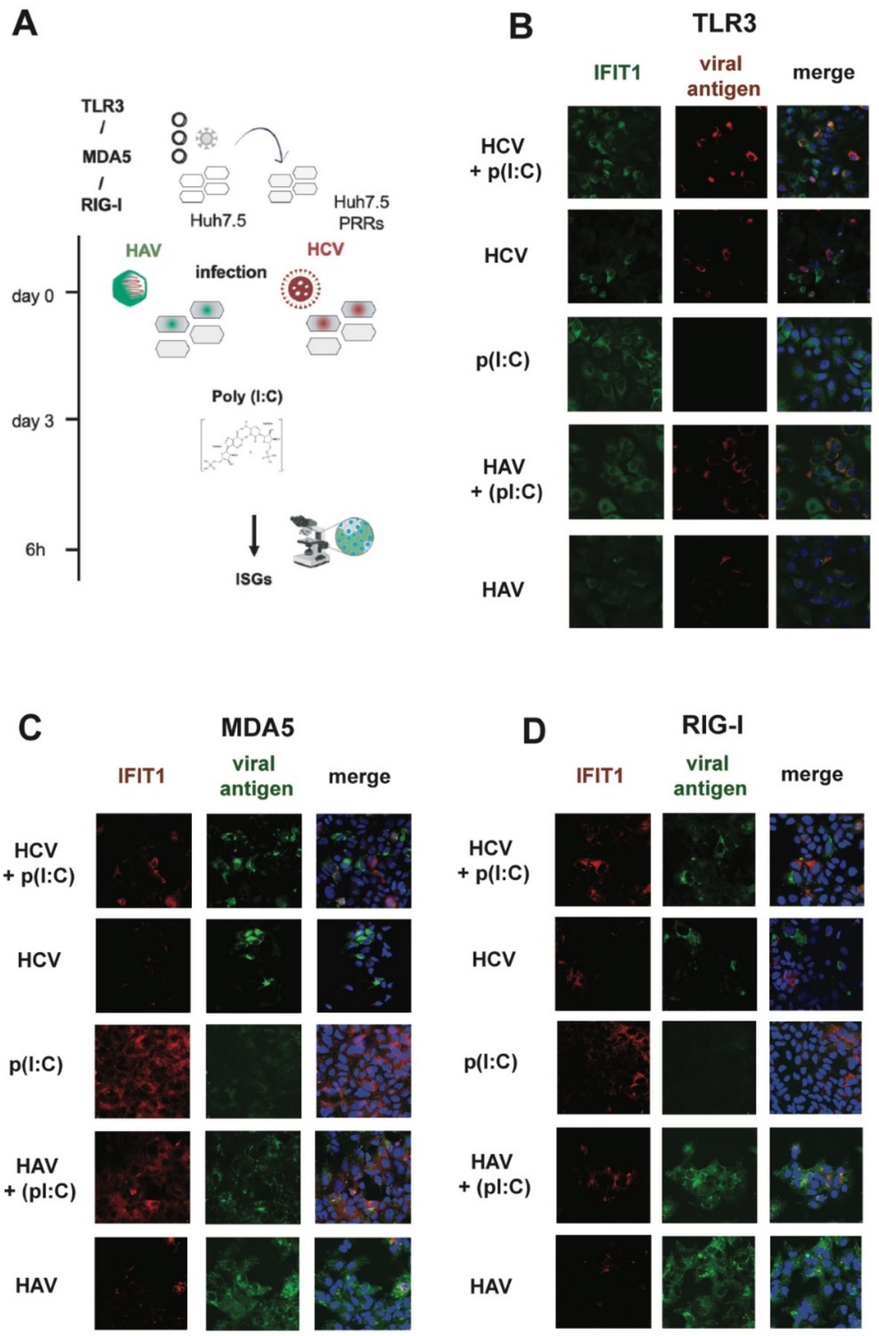


**Supplemental Figure 7. Interference with p(I:C) induced innate immune induction by HAV or HCV infection.**

(A) Schematic of the immunofluorence-based approach to assess counteraction of innate immune response in infected cells. Huh7.5 cells expressing TLR3, MDA5 or RIG-I were infected with HAV or HCV. On day 3 post infection cells were transfected with p(I:C), 6 hours later cells were fixed and stained for viral antigen and IFIT1 (ISG) expression. (B-D) Huh7.5 cells were transduced with lentiviral vectors encoding either TLR3, RIG-I or MDA5. After selection and passaging, cells were infected with either HAV (MOI 4) or HCV (MOI 1) and, after three days, seeded onto coverslips and transfected with p(I:C) or mock treated. After six hours, cells were fixed and stained for viral antigen (HAV 3C or HCV NS5A) as well as for IFIT1 with specific antibodies, nuclei were stained with DAPI.

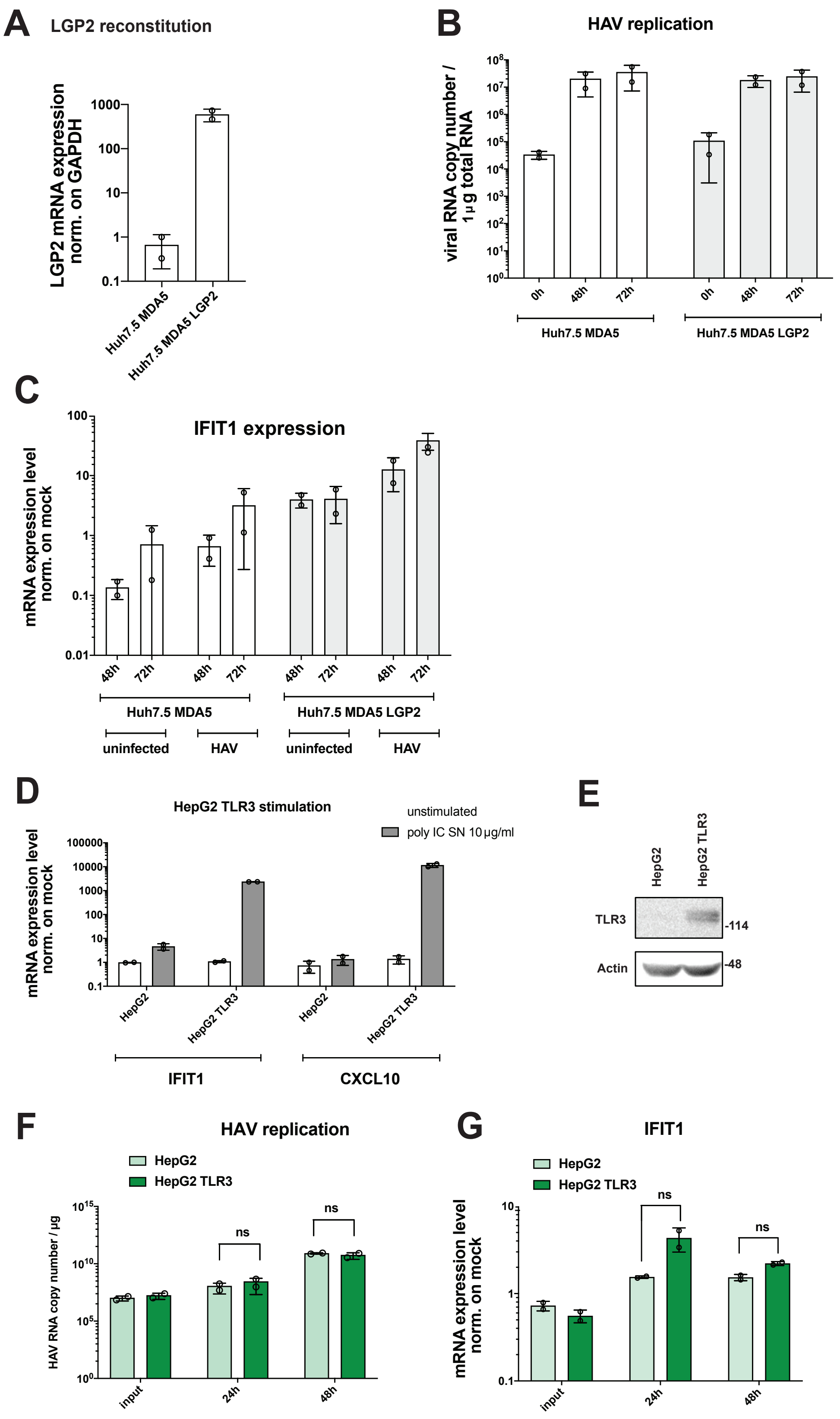


**Supplemental Figure 8. HAV mediated innate immune induction in HepG2 cells.**
(A) Huh7.5 cells stably expressing MDA5 were transduced with lentiviral vectors encoding LGP2. After selection and passaging, total RNA was isolated from the cells and LGP2 mRNA

levels were measured via RT qPCR and normalized to GAPDH expression. Data are shown as fold expression relative to Huh7.5 MDA5 cells without reconstituted LGP2. (B, C) Huh7.5 MDA5 cells, empty or stably expressing LGP2, were infected with HAV HM175/18f using a MOI of 4. At the indicated time points total RNA was isolated and viral RNA (B) along with IFIT1 mRNA expression levels (C) were measured. IFIT1 mRNA levels are shown as fold expression relative to uninfected MDA5 cells without reconstituted LGP2. All values shown are mean values with SD from biological replicates (n = 2).

(D) HepG2 cells were transduced with a lentiviral vector encoding TLR3. After selection and passaging, cells were subjected to p(I:C) supernatant feeding to exclusively address the TLR3 pathway. Six hours after stimulation, cells were harvested and total RNA was isolated; IFIT1 and CXCL10 mRNA expression levels were measured by RT-qPCR. (E) 10 µg of protein lysate were separated by SDS-PAGE and subjected to immunoblotting, using specific antibodies against TLR3 and b-actin (Actin), as indicated. (F-G) HepG2 cells with or without stable TLR3 expression were infected with HAV HM175/18f at a MOI of 4. At the indicated timepoints, cells were harvested and total RNA was isolated. HAV RNA (F) and IFIT1 mRNA expression levels (G) were quantified via RT-qPCR. IFIT1 mRNA levels are shown normalized to GAPDH expression. Data are shown as fold expression relative to uninfected cells. All values shown are mean values with SD from biological duplicates (n = 2).

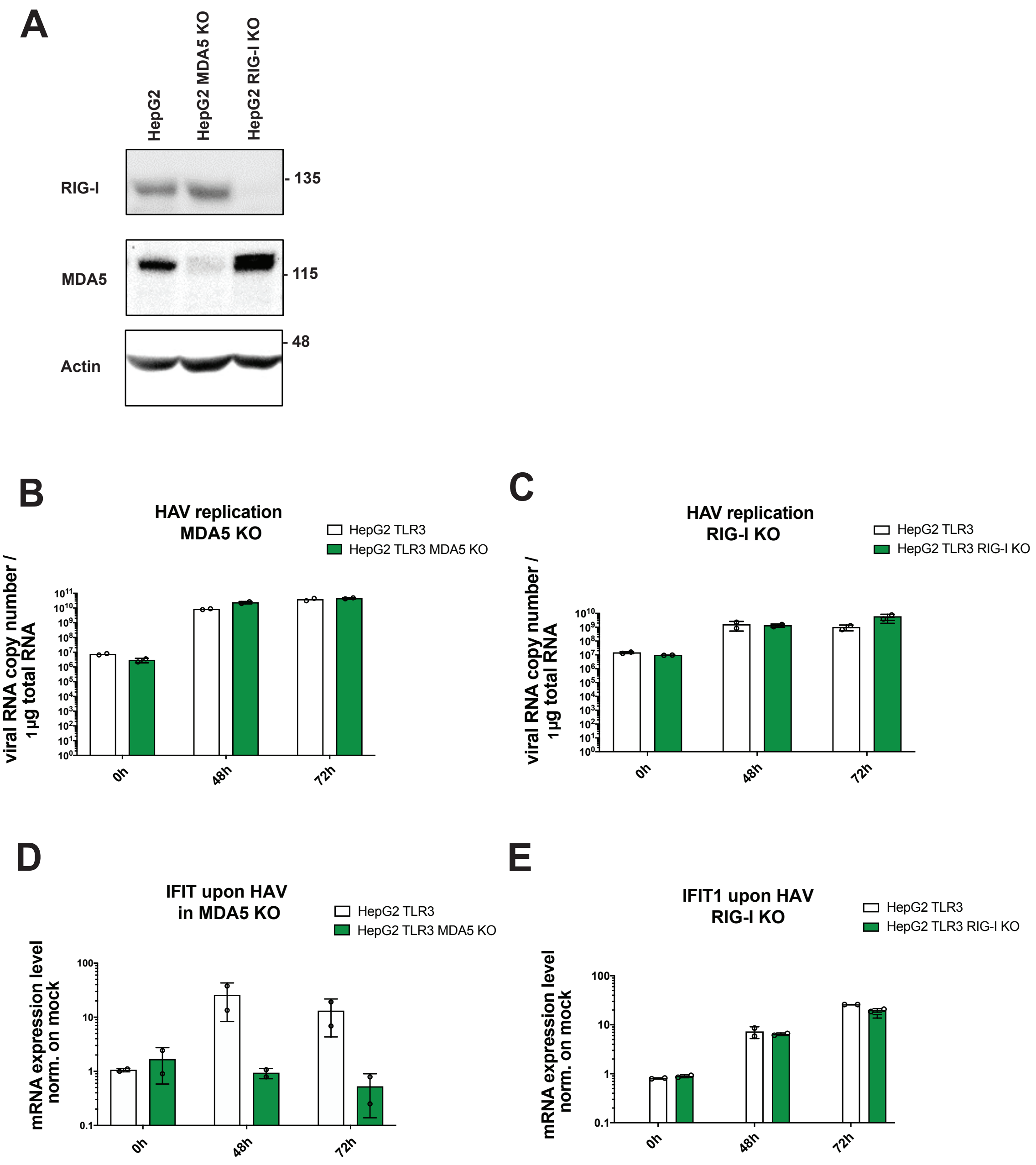


**Supplemental Figure 9. Determinants of HAV sensing in HepG2 and Huh7 cells.**
(A) HepG2 cells and cell pools with knockout (KO) of RIG-I or MDA5 were analyzed by immunoblotting, using specific antibodies against RIG-I, MDA5 and b-actin (Actin), as indicated. (B-E) HepG2 TLR3 cell pools with knockout (KO) of MDA5 (B, D) or RIG-I (C, E), respectively, were infected with HAV and analyzed for IFIT1 mRNA (D, E) and viral RNA (B, C), at the indicated time points. IFIT1 mRNA levels were normalized to GAPDH expression and shown as fold expression relative to uninfected cells. All values shown are mean values with SD from biological replicates (n = 2).
